# Markerless tracking of an entire insect colony

**DOI:** 10.1101/2020.03.26.007302

**Authors:** Katarzyna Bozek, Laetitia Hebert, Yoann Portugal, Greg J. Stephens

## Abstract

We present a comprehensive, computational method for tracking an entire colony of the honey bee *Apis mellifera* using high-resolution video on a natural honeycomb background. We adapt a convolutional neural network (CNN) segmentation architecture to automatically identify bee and brood cell positions, body orientations and within-cell states. We achieve high accuracy (~10% body width error in position, ~10° error in orientation, and true positive rate > 90%) and demonstrate months-long monitoring of sociometric colony fluctuations. We combine extracted positions with rich visual features of organism-centered images to track individuals over time and through challenging occluding events, recovering ~79% of bee trajectories from five observation hives over a span of 5 minutes. The resulting trajectories reveal important behaviors, including fast motion, comb-cell activity, and waggle dances. Our results provide new opportunities for the quantitative study of collective bee behavior and for advancing tracking techniques of crowded systems.

## Introduction

Among the rich phenomenology of organismal behavior, honey bees and other eusocial animals are distinguished by their remarkable, self-organizing, collective dynamics on the scale of an entire society. Functioning as a “super organism”, a honey bee colony can contain thousands of individuals whose intricate behavior results from a shared (though not clonal) genetic background and sophisticated social signals conveyed through multiple communication channels ^1^. The effect of these dynamics is to cooperatively divide and organize the effort necessary to maintain a well-functioning colony in response to external and internal environmental change, and to grow. A longstanding fascination with such behavior has driven substantial previous work (see e.g. ^2^) which includes examinations of collective behavior ^3,4^ also in combination with high-throughput sampling technologies such as gene expression sequencing ^5–7^. However, a full quantitative understanding of the colony behavior requires measuring the collective dynamics at single organism resolution as well as the spatiotemporal patterns of colony resources such as food and brood. Both of these challenges are now accessible through advances in machine vision.

A natural honey bee colony contains a high density of visually similar members, rapidly moving and occluding on the uneven and changing honeycomb surface, and whose numbers change in time. These factors present substantial difficulties for automated image analysis for which a common solution is to apply physical tags to some ^8^ or most ^9,10^ of the colony members. Barcoded tags allow for the distinct marking of a sufficiently large number of individuals to track a naturally-sized colony and have been exploited to unravel important aspects of bee communication ^8,11^ and information spread ^10^. However, the burden of manually tagging hundreds or thousands of small insects, without harm or inhibition to their motion, carries severe limitations. For example, marking newly hatched bees requires either opening the hive or introducing marked newborns without letting any hatch inside, both of which disrupt the colony ^9,11^. Moreover, the recognition of markers fails when the tag is partially occluded or blurred. In the dense environment of a hive, even when confined to a 2D surface, partial occlusions are common. In particular, behaviors such as crawling inside of a honeycomb cell, or walking upside down on the glass of the observation hive, can hide the marker, thus limiting behavioral repertoires captured with a marker-based approach. Physical tags cannot be used to identify brood, honey or pollen stores, and the difficulty and workload of manual tagging hinders the analysis of multiple colonies, as is routinely accomplished in behavioral studies of other organisms.

Recent breakthroughs in image analysis using CNNs, including fast and accurate single and multiple object detection ^12–14^, posture quantification ^15–18^ and image and video appearance representation ^19–21^, offer new inspiration and opportunities for extracting information directly from video data in dense-animal contexts. However, most existing solutions and benchmark datasets for multi-object tracking as well as for posture and activity recognition, are dedicated to human behavior and crowds ^22–25^. In biological image data CNNs have been broadly exploited for cell or particle segmentation ^26–29^. Versatile, supervised CNN-based tools have also been proposed for animal posture quantification ^17,18^ and have been successfully applied to the study of insect behavior ^30–32^. While facilitating important tasks in bioimage interpretation, these solutions are limited to behavior of few individuals and do not resolve challenges within the task of dense object detection and tracking in a bee colony. Importantly, previous work has shown that seemingly identical organisms do carry distinct visual features, which can be quantified and leveraged for markerless tracking ^33–35^.

Here we harness machine vision through CNNs to capture, at single-organism resolution, the colony-wide composition and behavior of the honey bee *Apis mellifera.* Our approach applies to images and video recordings of unmarked colonies and enables broad quantitative study without the burden of manual marking. We demonstrate our solution through the analysis of five long-term timelapse recordings, up to four months in duration (segmentation and sociometry), as well as of five short-time videos recorded at high frame rate for five-minutes (tracking and motion behavior). The data was collected at various locations on the campus of OIST Graduate University (Okinawa, Japan) with varying imaging arrangements. We infer the position, orientation and within-cell state of each bee together with the location of brood cells (Fig. 1A). Using these detection results we devised a neural network and an efficient training method for quantifying visual features capturing similarity among bee instances (Fig. 1B). We use this similarity to stitch bee detections into trajectories across difficult occlusion and touching events. We demonstrate our approach with long-term sociometric monitoring (detection) and with colony-wide exploration of individual behaviors (tracking). Along with this manuscript and associated code, our contribution includes the labeled image data of thousands of trajectories of bees in dense configurations, a unique resource which offers new opportunities for both machine vision and biological research.

**Figure 1.**
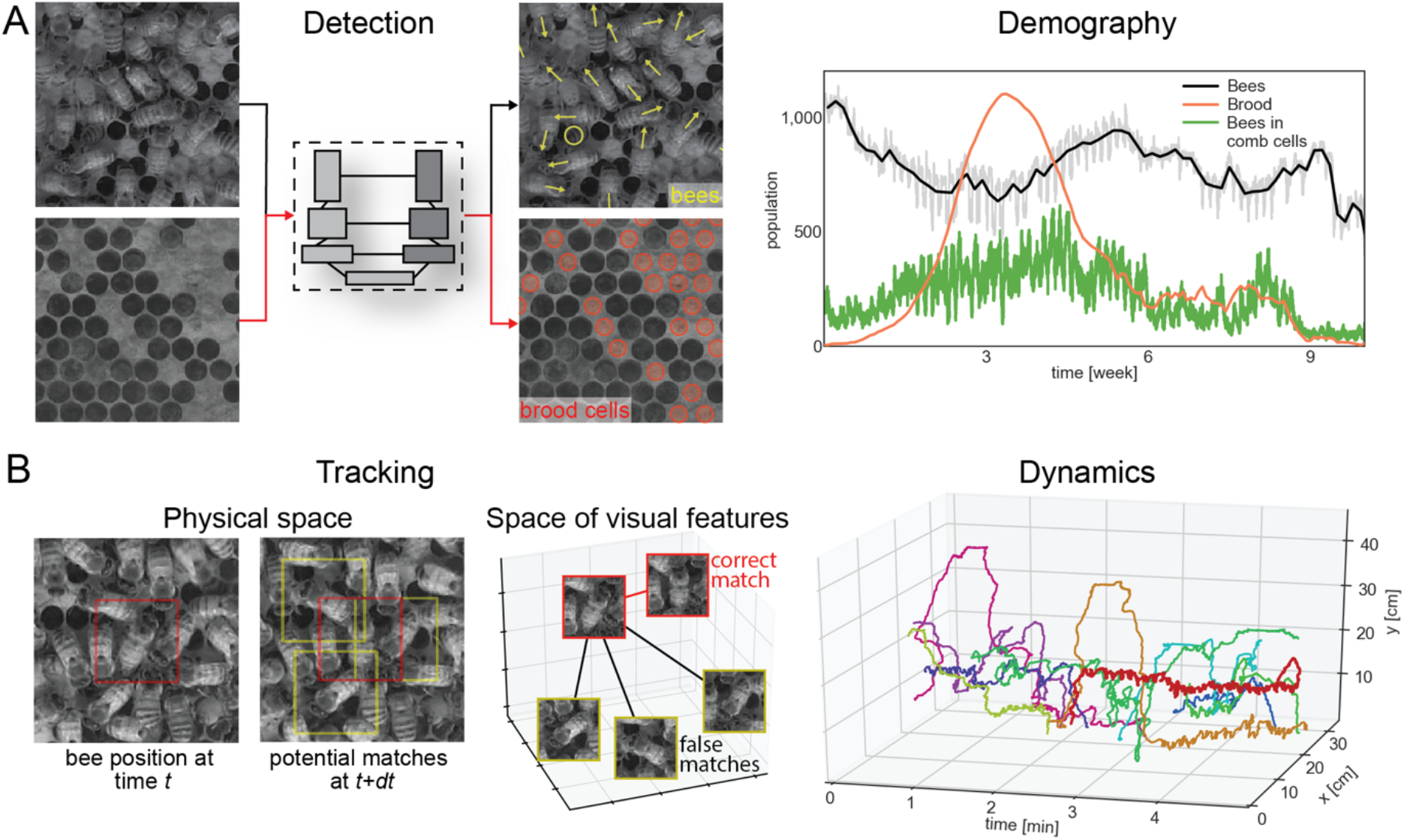
Schematic of the detection and tracking methods. **A.** A segmentation architecture was adapted for the detection of bees and brood cells in the dense environment of a bee colony. We use this architecture to infer bee and brood positions (yellow and red marks, respectively), bee posture type – bee inside of a comb cell (yellow round symbols, denoted later as cell-bees) and fully visible bee (yellow arrows, denoted later as full-bees) – and the orientation angle of the fully visible bees (angle of the yellow arrows). Bee and brood locations, as well as within-cell state, allow for detailed sociometric analyses over long timespans and across multiple colonies (right). **B.** CNN-derived embeddings of visual bee features are used to link detections across video frames into trajectories. The network is trained to maximize the Euclidean distance between detections belonging to different trajectories and minimize this distance between detections of the same bee within one trajectory. This visual similarity metric allows for accurate construction of trajectories of unmarked bees and analysis of dynamic aspects of bee colonies. (right) For illustration we show ten tracked trajectories, likely corresponding to forager bees. The red trajectory belongs to a bee that entered the hive around minute 2 of the recording and performed rapid back-and-forth dancing motion for the next 2 minutes.

## Results

### Long-term colony sociometry

To capture the long-term sociometric dynamics ^36–39^ of bee colonies, we devised a segmentation-based method for the detection of bees and brood in dense configurations within a 2D hive ^13,35^. For bee detection our solution is based on a modified segmentation CNN architecture ^12^ which exploits the temporal dimension of video data to improve accuracy (Fig. 2A-B). During training and inference this network uses information from the preceding video frame, allowing us to reduce the size of the network by 94% compared to the original architecture while still achieving high detection accuracy. Within one network we infer both the segmentation maps through pixel classification as well as the orientation angle of the segmented objects through regression. The segmentation maps delineate only the central parts of the bees’ bodies or central parts of the abdomens of bees inside of the comb cells. Each pixel is classified into one of the three categories: ‘background’, ‘fully visible bee’ (denoted full-bees throughout the text), and ‘bee inside comb cell’ (denoted cell-bees throughout the text) if only the bee abdomen is visible. The position of each bee is inferred as the central point of the respective foreground pixel group. We infer an orientation angle only for the full-bee class. The resulting segmentation maps allowed to find individual locations (Fig. 2C) in an independent test set with a precision of < 10% of a bee body width, detection true positive rate (TPR) ~ 0.96, false positive rate (FPR) ~ 0.14 and orientation angle error of ~ 9.7° closely matching error of human labelers (Supplemental Table T1). The segmentation results above were first reported in an earlier conference proceedings^13^.

**Figure 2.**
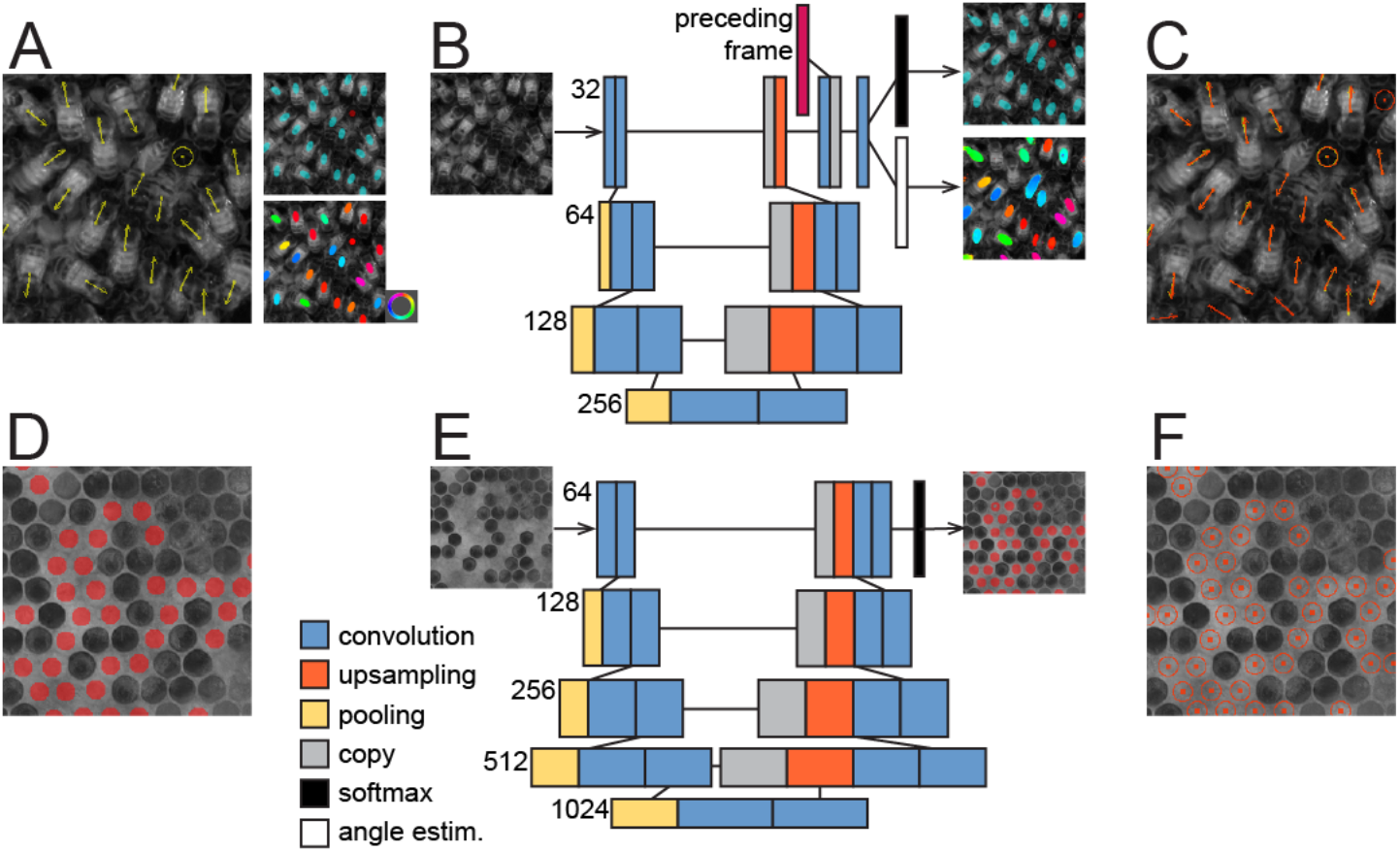
Dense object detection. **A.** Our manual annotation consists of the central points of each body, object class – full-bees (yellow arrows) or cell-bees (yellow round symbol) – and the body orientation of full-bees (arrows). Segmentation maps are created with foreground pixels denoting central parts of bee bodies. The upper segmentation map indicates class information, the bottom segmentation map indicates the orientation angle (color wheel). **B.** We modified the U-Net architecture^12^ by including a recurrent component which exploits temporal information from the preceding video frame in the penultimate network layer. The recurrent component allowed us to reduce the number of parameters by ~94% from the original U-Net (shown in E) without compromising accuracy. We added two loss functions to the network, one for class and one for orientation angle estimation, and their example output is shown to the right. **C.** Position, class, and orientation information is inferred from the network output. Red markers indicate inferred detections, labels are in yellow. **D.** Segmentation maps created for the training of brood cell detection. Similar to body detections, the foreground pixels cover only the central parts of the cells, allowing for object counting and localization. **E.** The original U-Net architecture is used for brood cell detection. **F.** Inferred brood cell positions.

The accuracy of the detection method was assessed on a large set of manually labeled images. We additionally inspected its performance in three frames from recordings L5 and S5 after retraining on a set of up to five frames from recordings L1-L4 and S1-S4. In frames from the beginning, middle, and end parts of recordings L5 and S5 that were not included in the retraining, we estimated detection FPR at < 1.4 % and false negative rate (FNR) at < 1% resulting in precision and recall >= 0.98 in these recordings (Supplemental Fig. S1). These results confirm the capacity of our detection algorithm to generalize across recordings and also provide a strong foundation for further analysis and tracking method development.

We devised a similar segmentation-based approach to locate brood cells. Due to the low temporal resolution of the background images generated for time spans of 12 h (Supplemental Movie M1), this solution did not include the recurrent element in the network design and relied on the original segmentation architecture ^12^ (Fig. 2D-E). We used round-shaped brood cell segmentation markers (Fig. 2F, Supplemental Fig. S2, Supplemental Movie M2) with no overlap and achieved a detection accuracy of TPR ~ 0.95, FPR ~ 0.02, FNR ~ 0.05.

We deployed both detection methods on a set of long-time video recordings (Supplemental Table T2) and extracted quantitative measures of demographic changes in bee colonies over periods ranging from two weeks to four months. In all long-term recordings (L1-L5) we found that the total population (full-bees and cell-bees together) in each colony as well as the proportion of cell-bees undergo repetitive fluctuations (Fig. 3A-B, Supplemental Fig. S4-5). The period of these fluctuations is close to 24 h based on spectral analysis (Fig. 3A-B, Supplemental Fig. S6). Over longer times, peaks in population followed peaks in brood numbers (Fig. 3C) and we found a strong negative correlation between fluctuations in population and brood counts (Fig. 3D, mean R^2^ < −0.84, P <0.0001 over the first three weeks), perhaps an indication of homeostatic control of colony size.

**Figure 3.**
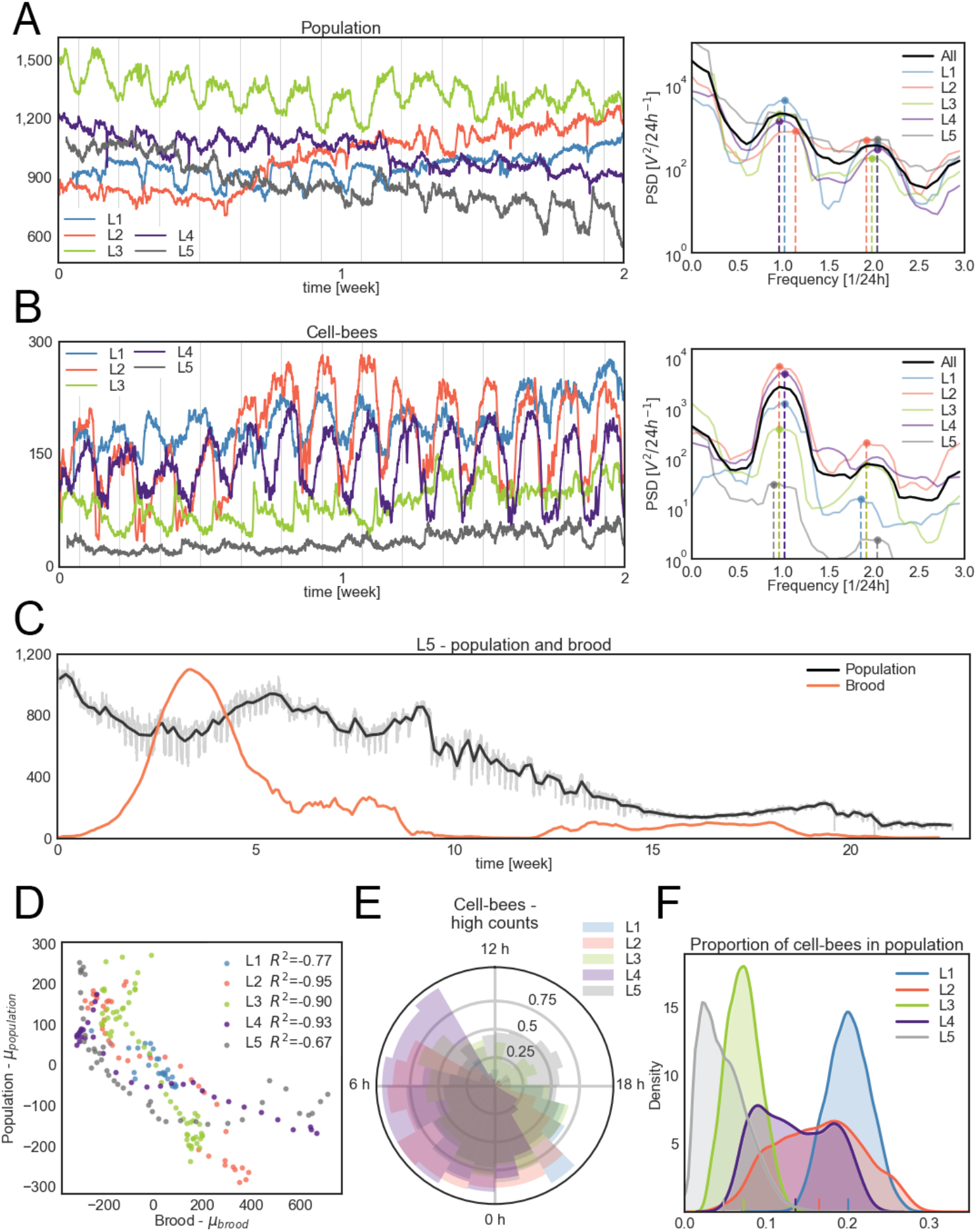
Colony sociometry. **A.** Daily fluctuations of the total population (full-bees + cell-bees) in hives L1-L5. The period of the fluctuations is approximately 24 h in all hives which is indicated by the power spectral density (right). Vertical lines denote midnight. **B.** Analogous to (A), daily fluctuations in the numbers of cell-bees and the respective power spectral density. **C.** Total population (full bees + cell-bees, gray and black lines) and brood population (orange line) in bee colony L5. Colony L5 was imaged for over four months and exhibited a population decline and ultimately colony collapse. After an initial rise in brood population, another increase of such amplitude did not occur, and the population steadily declined from week 10. The black line indicates the daily average population size while the exact count in each image is shown in gray. **D.** Fluctuations in bee and brood populations are anti-correlated. Total population averaged over consecutive 12 h windows is plotted against the brood counts in the same time window in the 5 colonies for the period of up to three weeks. For each time series, the mean of the series was subtracted from the plotted values. **E.** The nightly presence of cell-bees. We divide each 24 h of a recording into 24 1-hour bins and count how often the number of cell-bees is above the daily median number. High numbers of cell-bees occur predominantly in the evening, between 21 h and 6 h. **F.** Distribution of the proportion of cell-bees relative to the total population in hives L1-L5. Hive L3 and L5 show markedly lower proportion of cell-bees, which may be related to colony declines during our recording.

We next investigated daily changes in the number of full-bees and cell-bees as potentially reflecting the daily foraging, brood nursing, and other activities in each colony. We found high numbers of cell-bees predominantly at night (Fig. 3E) which may represent sleeping bees ^40^. This regularity was not apparent in the full population, where the oscillations varied in their phase, perhaps due to external factors (Supplemental Fig. S7). We thus inspected the weather conditions during the time of recordings and, while no extreme events were observed (Supplemental Fig. S8), two colonies (L3 and L4) were filmed during particularly high temperatures exceeding 26°C at night, which can encourage bees to stay at the hive entrance ^41^ where they would not be detected.

While we found insufficient evidence linking the phase of the population size change to the weather conditions, we did find that full-bee numbers shift in relationship to brood presence. During times when brood numbers are high, the colony population is in phase with the cell-bee population (Supplemental Fig. S9) with high numbers at night. Such a pattern potentially corresponds to the foraging activity of the bees performed uniquely during the day ^2^ and it is more prominent in hives with high brood numbers which might reflect the increased food demand of growing colonies (Supplemental Fig. S9).

Similarly, the cell-bee population may be related to brood presence. In three of the five long-term recordings the proportion of cell-bees was positively correlated with the brood count (L3-L5, R^2^ > 0.4, p < 0.05). Additionally, their phase relationship suggests thermoregulatory activities aimed at maintaining constant brood temperature at night. We also found that the spatial distance between cell-bees and the brood cells exhibits a 24 h periodicity (Supplemental Figures S10-S11). The cell-bees are significantly closer to the brood cells than other bees (Wilcoxon test p < 0.0001 in L1-L4) and this distance decreases during the night and increases during the day.

In two of the long-term recordings – L3 and L5 – the colonies experienced a systematic decrease in the number of individuals. In L3 we noticed a moth infestation in the wax towards the end of the recording (Supplemental Fig. S12) which might have weakened the colony. In recording L5 each subsequent increase of the brood and colony size was lower than the previous one (Fig. 3C) ultimately leading to the gradual depletion of the colony. Despite an initial rapid increase in brood numbers, another wave of brood of such size did not occur and the colony never recovered its initial population size. In hives L3 and L5 the proportion of cell-bees was noticeably lower (Fig. 3F). The cell-bees in those hives were additionally located much farther away from the brood cells than in the healthy colonies (Supplemental Figures S10-S11).

While the specific reasons for the decline of these colonies are not clear, we show quantitative measures indicative of behavioral changes in unhealthy bee colonies. Indeed, the colony collapse phenomenon currently threatening bee populations worldwide have been ascribed to a range of factors ^42,43^. However, despite a range of theoretical models ^44,45^, no quantitative, data-driven assessment has yet been proposed. Unraveling the exact reasons for and mechanism of honey bee colonies collapse would require multiple comparative observations across a range of conditions. Our approach enables the study of this phenomenon in a time-resolved, comprehensive, and quantitative manner.

### Colony-wide tracking at single-organism resolution

Our high detection accuracy provides a strong foundation for reliably following objects in time. Even though the observation hive is designed to constrain a colony to a 2D surface, in practice many bee activities occur in 3D and occlusions are common. Events such as hiding inside of a comb cell, walking ‘upside down’ on the glass covering the hive, or walking on uneven parts of the comb surface, all increase the difficulty of automated tracking by creating occlusions leading to identity swaps. To minimize the effects of the 3D events on trajectory construction we devised a trajectory linking method incorporating not only detection coordinates but also class and posture information together with a quantitative encoding of visual features of individual bees.

Previous studies ^34,35,46^ have demonstrated that objects indistinguishable to human eye might nonetheless carry unique visual signatures, also termed ‘pixel personality’, that can be quantified and can importantly aid the task of simultaneous tracking of multiple highly similar objects. Notably, the rise of deep learning has enabled new ways of extracting such visual signatures from images. However, using CNNs for the quantification of pixel identities requires a set of instances of each object that can serve as a training set. For example, in tracking up to 100 fish such a set could be constructed if a segment in a video existed where no fish collisions were observed during 3,000 video frames ^46^. Notably in the much denser environment of ~1,000 individuals inside of a beehive, such collision-free segments are rare or nonexistent. A previously proposed solution ^35^ therefore extracted pixel identities of bees based on a smaller number of instances, re-learning these identities after each trajectory matching step. Unfortunately, this solution offered only limited accuracy and came with a high computational cost, compromising the reproducibility of the results.

Here we present a method based on a quantitative representation of visual features of the tracked objects, instead of their distinct pixel fingerprints. Quantitative visual features are extracted from a CNN^47^ trained using a triplet loss objective function ^48,49^ – a function which enables the expression of similarity among entities via their vector embeddings. During training, triplets of images are fed into the network and we search for a solution where the Euclidian distance between vector embeddings of two images of the same object at different time points is as small as possible while the distance between vector embeddings of two different objects is large (Fig. 1B). The major hurdle in training such a network for bee tracking is the massive space of all possible triplets containing two instances pertaining to the same trajectory and a third instance pertaining to any other time point of any other trajectory. To address this problem and enable network loss function convergence in a reasonable time, we devised a dedicated training scheme. Briefly, image triplets were sampled from training set trajectories that included bee detections neighboring in time and space which are sources of potential identity swaps. Additionally, the embeddings of all triplets used during training were simultaneously tested for correctness and the incorrect ones – where the distance criteria between positive and negative matches were not met – were used again in the network training.

The network was trained on a large, incrementally constructed set of validated bee trajectories. We used four 5 min-long recordings (S1-S4) captured in different locations on the campus with cameras of varying pixel resolutions (Supplemental Table T2). While the hives were all the same spatial size, the colonies varied in number of organisms, ranging from N=805 to N=1,316, and in their dynamics (Figures 4A, 5A). All videos were recorded during daytime, under good weather conditions, and within the foraging season of Okinawa.

**Figure 4.**
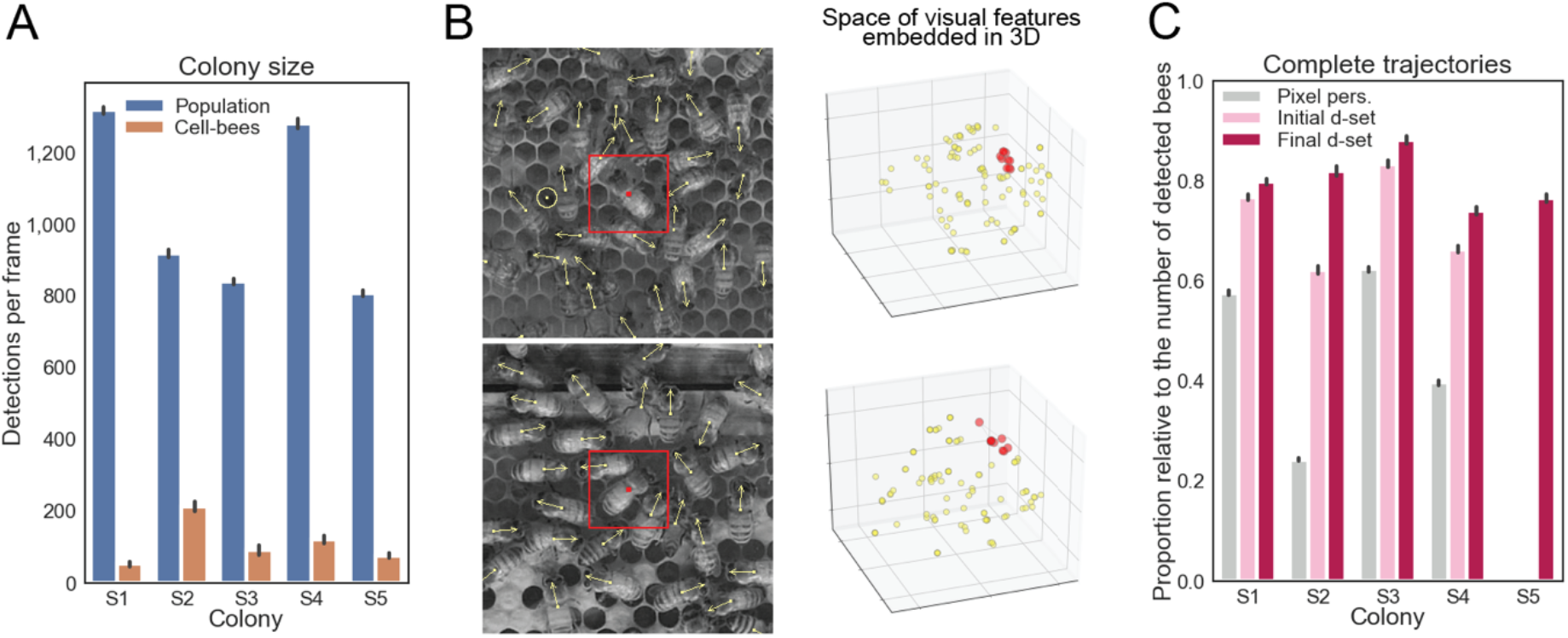
Leveraging visual features for enhanced detection matching across frames. **A.** Recordings from five different beehives were used for tracking method development and testing. The hives contained varying total populations and of bees in comb cells (cell-bees). **B.** CNN-derived vector embeddings are used to better match bee detections across video frames. The embeddings encode similarity among detections. Originally 64-dimensional, an example projection of the embeddings in 3D is shown in the panels on the right obtained with the use of t-SNE ^75^. Red dots represent embeddings in 10 consecutive video frames of the bees marked by red squares in the left panels. Embeddings of other bee detections in these images that occurred over consecutive 3 video frames are marked by yellow dots in the panel on the right. Even though bees appear identical to a human observer, the embeddings belonging to one trajectory (red) are distant from embeddings of all other bees neighboring in time and space, thus aiding the correct stitching of individual detections. **C.** The accuracy of the tracking method obtained for test recording S5 matched the accuracy reached in recordings S1-S4 used for network training and method design. We show the proportion of extracted trajectories relative to the average detected number of bees in hives S1-S5. The proportion of extracted trajectories is shown for an earlier approach (`pixel personality’, gray bars), the current method but trained only the initial dataset (pink bars) and for the current method but trained on the final dataset (red bars).

**Figure 5.**
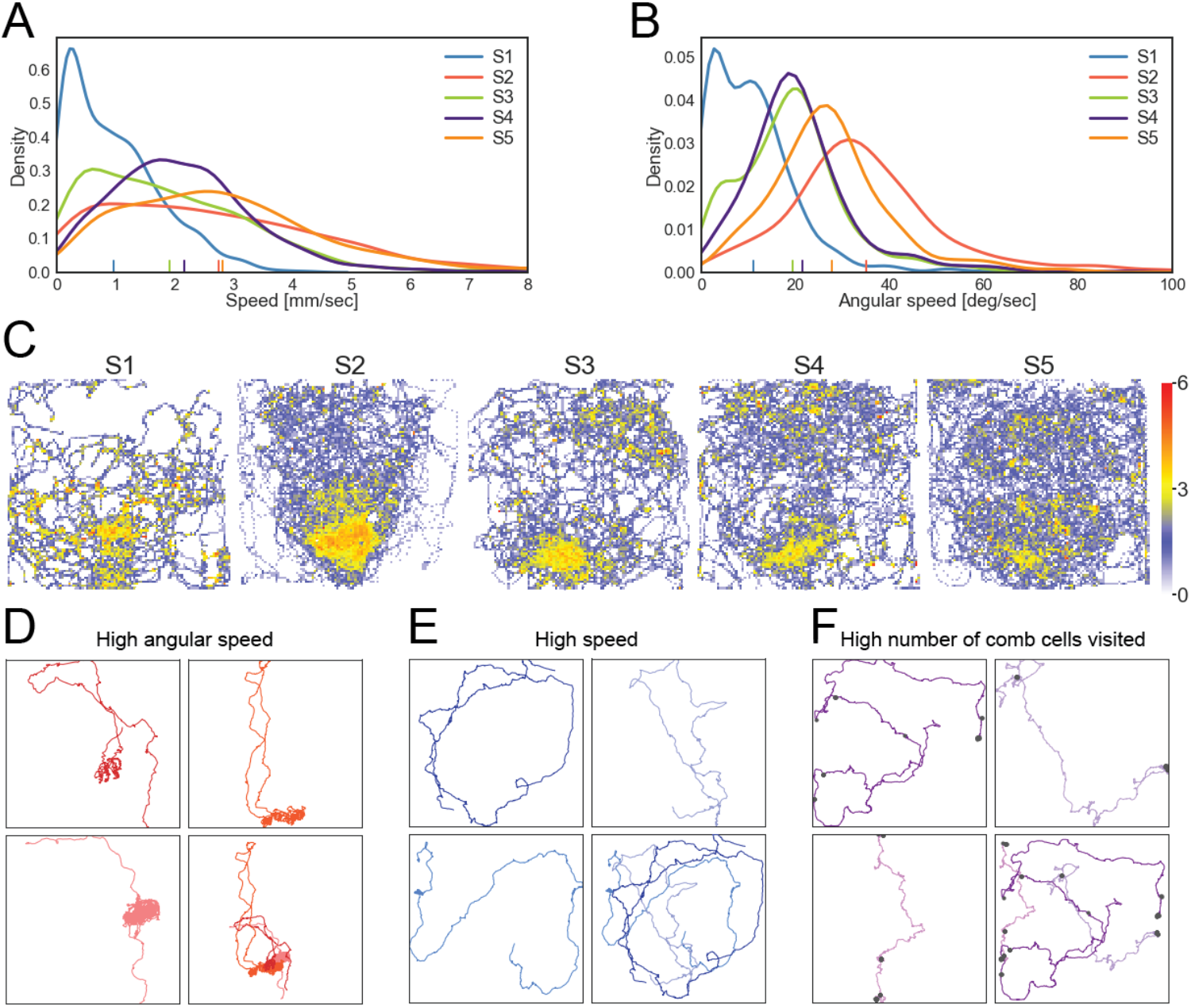
Colony dynamics at single-organism resolution. **A.** The distribution of individual mean speed computed from trajectories in hives S1-S5. Large differences across hives include a significant proportion of immobile bees in S1 and increased number of fast-moving bees in S2 and S5. **B.** The distribution of individual mean angular speed of trajectories in hives S1-S5. Low angular speeds are seen primarily in S1 while the largest proportion of trajectories with high angular speed is present in S2. Trajectories in the tail of these distributions are excellent candidates for forager bees performing or following a waggle dance. **C.** The spatial distribution of trajectories characterized by large linear and angular motion. We show 100 trajectories from each hive which have large linear and angular speed. Across all hives these trajectories are located near the entrance, which is at the bottom of each hive. This localization may reflect a forager recruitment site. **D.** Example trajectories of individuals with large linear and angular motion. Three trajectories are plotted individually and combined in the bottom-right panel. The densely overlapping parts of these trajectories indicate the location of the waggle dance performed by these individuals. **E.** Example trajectories showing large linear but low angular motion. Such individuals tend to move rapidly over large portions of the hive. **F.** Example trajectories of bees exhibiting a large number of comb-cell visits.

We first applied a “pixel personality” approach ^35^ on recordings S1-S4. The trajectories resulting from this approach were manually validated, and the correct trajectories formed the *’initial dataset’* for the learning of visual features’ embeddings as described above. Visual feature embeddings extracted from the CNN incorporate similarity among individuals (Fig. 4B, Supplemental Movies M3-M7) and hence allow the linking of bee detections across video frames in presence of temporary object occlusions. We implemented a detection matching solution exploiting both position and visual features, and matchings are done in a greedy manner on a sorted list of all pairs satisfying predefined time and space proximity criteria (see Methods).

Embedding-based trajectories in videos S1-S4 were next manually validated. Compared to the pixel identity approach, our solution offered an important increase in accuracy at a low computational cost (Fig. 4C). Visual feature-based matching resulted in a higher number of correct trajectories over the pixel personality approach as well as over a position only-based solution (Fig. 4C, Supplemental Figures S13-S14). Even though some trajectories belonging to these recordings were part of the training set, this result suggests the method’s capacity to generalize to other, trajectories not included in the train set within the same colonies.

Next, we used the entire set of trajectories validated as correct in videos S1-S4 to form the *‘final dataset’*. Embeddings derived from the network trained on the expanded training set delivered an increase in tracking performance in videos S1-S4 (Fig. 4C), emphasizing the role of the training set size in producing more precise and distinct embeddings. Importantly, we also tested a range of data augmentation scenarios as well as background masking (Supplemental Fig. S3) that did not positively affect tracking accuracy (Supplemental Fig. S15). We attribute this to the fact that data augmentation importantly increases data complexity hindering network convergence. Poorer results on images not including background additionally suggest that it plays an important role in allowing for correct matching.

To test the capacity of our method to generalize to other recordings, we used the recording S5. No images of this colony were used in the training of the detection or tracking method. Recording S5 is additionally characterized by vibrations due to neighboring construction site and flickering of the lights, creating particularly challenging tracking conditions. We found that 77% of bees were correctly tracked (Fig. 4C), a result comparable to the recordings that were part of the training set (Supplemental Table T3). Such high accuracy provides a strong foundation for the study of colony dynamics and we expect improvements as more validated trajectories are incorporated in the training set. The entire set of correct trajectories assembled during development of this method includes 4,642 trajectories from the short-term recordings S1-S5 and is provided with this manuscript as an important resource for further study and method development.

### Quantitative analysis of colony behavior

The large set of trajectories generated through our detection and tracking approach covers an extensive proportion of the recorded colonies (Supplemental Movies M9-M13) enabling a broad, comparative study of honey bee colony dynamics. While our emphasis here has been on the techniques of detection and tracking, we can already offer a variety of quantitative observations.

We first compared aggregate dynamics using the distributions of individuals’ speed, angular speed, diffusion coefficient, as well as motion span defined as the diagonal of the minimal rectangle fitting the trajectory (Fig. 5A-B, Supplemental Fig. S16). Colonies S1 and S3, despite differing in the number of individuals (1,316 and 839, respectively), are characterized by a large proportion of motionless individuals and individuals moving over only limited areas of the hive (8% of the colony diagonal on average in S1). Whereas colonies S2 and S5 showed a high proportion of fast-moving bees and bees traversing larger portions of the surface of the hive (~24% of the high diagonal on average). Additionally, colonies S2 and S5 contained a higher proportion of individuals moving at high angular speed (Fig. 5B) which could be indicative of the foragers performing waggle dance ^1^.

We next inspected the spatial localization of the individuals showing the highest values in each motion attribute (speed, angular speed, area of motion, etc.). Most prominently, bees showing the highest angular speed tend to be located at the bottom part of the hive in proximity of its entrance (Fig. 5C). The colocalization of these individuals, most noticeable in S3, is present across all colonies, suggesting that relatively high angular speed is a characteristic of foragers performing and following the waggle dance at the hive entrance ^50^.

To corroborate this hypothesis, we inspected trajectories showing high mean values of single motion attributes as well as their combinations. Trajectories with high angular and translational motion indeed belong to bees performing waggle dance (Fig. 5D). The high angular speed of these individuals is displayed during the fast looping dance motion of the foragers indicating to other colony members locations of food sources (Supplemental Movies M14-M16). In contrast, individuals with high translational motion but low angular velocity tend to visit large portions of the hive without performing any recognizable action (Fig. 5E), a potential sign of patrolling behavior ^51^. Lastly, we quantified the number of times a bee in each trajectory visits a comb cell and identified bees that clean or search through comb cells sometimes across long distances (Fig. 5F, Supplemental Movies. M17-M19). These examples demonstrate that the collected metrics group together similar and potentially meaningful individual bee behaviors.

## Discussion

Recent machine vision advances in the precise posture tracking of individual animals ^17,18,52^ as well as of the positions of highly-similar organisms in groups ^46^ are enabling new quantitative studies of behavior ^53,54^. In collective behavior specifically, the use of CNNs for the pixel-based identification of individual organisms has significantly advanced markerless, long-time tracking in 2D, from more modest assemblies (~10 individuals) ^34^ to large groups (~100 individuals) ^46^. However, while network-learned identities can resolve confounding occlusions and overlaps, a principal challenge of individual-resolution group tracking, there must also be enough isolated instances to train the identification network. These conditions do not exist in the dense and cluttered setting of a honey bee colony, where occlusions and overlaps are perpetual. Here we have described a detection-to-dynamics solution which expands behavioral tracking to large and dense collective systems.

Our markerless detection and tracking techniques offer new possibilities for the quantitative study of honey bee colonies on the collective scale at single-organism resolution and are complementary to existing approaches ^11^. Both the brood and adult populations change over time (Fig. 3), a challenge for manual marking, and (with improved lighting of the observation hive) our detection technique can be readily extended to include capped and uncapped honey as well as pollen cells. The dynamics and spatial arrangement of these variables provide new quantitative data for sociometric analysis ^36–39^ and will be particularly interesting in the context of a collapsing colony. Colony-wide, high-resolution tracking augments larger-scale measures such as weight ^55^ and can be combined with additional hive sensors for a novel surveillance system ^56^. The automatic nature of our approach also facilitates the imaging of multiple hives ^57^, an important consideration due to colony-to-colony variability.

Our trajectory construction method currently spans over five minutes, an interval chosen to enable extensive manual validation of the results (Fig. 4). We expect that this interval can be increased with more training data (trajectories from more colonies), advanced validation techniques (such as using markers invisible in the infrared), as well as with improved matching procedures. In particular, learning motion patterns via a recurrent neural network ^58^ can improve matching accuracy over the heuristic approach proposed here. Combining motion and appearance learning, while still underexplored ^59^, has potential to result in a single and powerful tracking solution. Indeed, an important outcome of this work is to provide first techniques together with rich trajectory data for further improvement as machine vision methods accelerate.

A wealth of behavioral information is already accessible within our current tracking window. While not a target of our study, we could readily detect bees performing the waggle dance based on their motion (Fig. 5B). Other possibilities include detection of trophallaxis ^60^, fanning and scenting ^61^, for which additional appearance cues can be used, such as the extension of the proboscis and wings. Additionally, the quantitation of behavior and aggregate colony dynamics over short time scales could be used in combination with the long term sociometric observation of the same colonies. The accuracy of CNNs in detection of visual detail together with the large numbers of trajectories of unmarked bees offered by our methods present vast new opportunities for behavior analysis of bees and open avenues for more quantitative approaches to modeling colony dynamics ^62–66^. We also see no obvious obstacles in generalizing our approach to other dense insect collectives such as ants.

Across the organizational scales of living systems, from molecular, through neural ^67^, to animal groups and societies ^68,69^, including humans ^70^, our ability to understand emergent collective behavior has been significantly enhanced by modern precision measurements and analysis across large portions of the ensembles. With the advances reported here, we expect accelerated progress in our understanding of the behavior of honey bee colonies and other crowded systems.

## Methods

We provide code, tutorials, and links to our honey bee datasets:

dataset: https://groups.oist.jp/bptu/honeybee-tracking-dataset#tra
code and tutorial: https://github.com/kasiabozek/bee_tracking
segmentation labeling: https://github.com/oist/DenseObjectAnnotation

### Imaging setup

We situated observation hives in two distinct locations: location 1 on the rooftop of an OIST building (approximately 4^th^-floor elevation), and location 2 in a ground-level shipping container which was also surrounded by greenery. Both locations were equipped with infrared LEDs and a heating system which maintained a constant room temperature of 31°C. The observation hives were 47 × 47 cm in size, fitting two honeycomb frames placed one above the other. The back side of the comb was fixed to a wooden surface constraining the bees to only one side of the frames. In location 1, images were obtained with a 5,120 × 5,120 pixel Vieworks Industrial Camera VC-25MX-M72D0-DIN-FM, at 30 FPS. In location 2, images were obtained with two lower resolution cameras Panasonic Lumix GH5 4K and Blackmagic Design Production Camera 4K at 30 FPS with a typical 4k resolution of 3,840 × 2,160 or 2,560 × 2,560 pixel for the long timespan recordings (Supplemental Fig. S1). Images of recordings from location 1 (S1, S2) were spatially down-sampled by a factor of two resulting in a similar pixel-per-bee resolution for the recordings from both locations. All cameras were modified for infrared imaging by removing the infrared filter.

### Detection dataset

For the development of our detection method we generated two recordings in location 1 (D1, D2) with spatial resolution of 5,120 × 5,120 pixel. Two sequences of 360 images from each of these recordings, spaced in time by 0.5 sec, were used for training and testing. We devised a custom labelling interface (https://github.com/oist/DenseObjectAnnotation) for manual annotation of bee locations and orientations. Through the interface the user defines a position and orientation of a fully visible bee (full-bee) by dragging, dropping and rotating an arrow symbol in an image. An additional round symbol with no orientation information was used to mark the abdomens of bees partially hidden inside a comb cell (cell-bees).

We used our interface to generate a labeled dataset through Amazon Mechanical Turk (AMT). We selected regions of size 3,072 × 2,048 pixel and of 3,072 × 3,072 pixel in images from recordings D1 and D2 respectively containing most of the colony bees against various backgrounds. These image regions were then submitted for annotation. We also submitted a subset of four frames – two from each recording – with 2,034 bees for labeling 10-times by independent workers to obtain an estimate of human error in position and orientation labeling. This error was calculated as the standard deviation of distance of each of the 10 labels to the reference label obtained in the main labeling task.

As a result of the labeling we obtained a dataset containing 375,698 labeled bees. Every bee in this set was assigned (*x*, *y*, *b*, *α*) denoting the coordinates of the central point of a bee against the top-left corner of the image, type of the label (b=1 for full-bees and b=2 for cell-bees), and the body rotation angle *α* against the vertical pointing upwards and calculated clockwise (*α* = 0 *if b* = 2). To use this information for segmentation-based individual localization we generated regions centered over the central point of each bee. For full-bees the regions were ellipse-shaped with axes r1=20 pixels and r2=35 pixels, roughly a third of the bee dimensions in the image. For cell-bees the regions were circular with r=20 pixels. Such regions cover central parts of each bee and are non-adjacent to regions covering neighboring bees in the image.

These foreground regions were assigned values of class b in the classification segmentation maps. Background pixels were assigned value 0. For learning of the orientation angles, each foreground pixel, instead of class label, was set at the value of the bee rotation angle and the background pixels were labeled as −1.

To compensate for the class imbalance between foreground bee regions and the non-bee background, we generated weights used for balancing the loss function at every pixel. For every bee region a 2D Gaussian of the same shape was generated, centered over the bee central point, and scaled by proportion in the training set of the background pixels to the number of bee-region pixels.

For training and testing the images were organized in 60 sequences of 360 images of 512 × 512 pixel size. In this time-resolved data the first 324 images of each sequence were used for training and the remaining 36 for testing.

### Detection network and training

We used the U-Net segmentation network^12^ and expanded its functionality to take advantage of regularities in the image time series patterns. In each pass of the network training or prediction the penultimate layer was kept as a prior for the next pass of the network (Fig. 2B). In the following pass the next image in the sequence was used as input and the penultimate layer was concatenated with the prior representation before calculating network output.

We used two loss functions in the last network layers. The first loss function, the 3-class softmax function, allowed for performing foreground-background segmentation. The second loss function was defined as 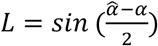 where 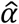 and *α* are the predicted and labeled orientation angle respectively.

In the network output, each contiguous foreground region was interpreted as an individual bee. Foreground patches smaller than 10 and larger than 1,000 pixels were discarded as potentially wrong. The centroid location was calculated as the mid-point of all x and y coordinates of points in each region. Region class was assigned as the class identity of the majority of pixels within given region. We also calculated the main body axis full-bee regions as the angle of the first principal component of the points in each region. The predicted orientation angle of the pixels within a given regions was used to assign back and front to the region principal axis. Due to the higher prediction errors in the image margins, where objects are not fully visible, we discarded results within a 50 pixel margin of the image boundaries.

### Recordings

We performed long-term time-lapse imaging of five colonies (L1-L5) for a timespan between two weeks to four months (Supplemental Table T2) with sampling frequency of once per minute (colony L5) and once every two minutes (colonies L1-L4). We additionally imaged five colonies (S1-S5) with a sampling frequency of 30 FPS. For the tracking analysis we used five-minutes segments, which we down-sampled to 10 FPS.

All image data was collected from spring to fall in favorable weather conditions when foraging activity was observable. For each recording we typically selected from the hives in our apiary two honeycomb frames with even surface, one containing brood and one containing food stores. We removed comb cells and other content on one side of each frame and left them for another day inside of the apiary hive to allow the bees to clean the damaged surface. Next, after ensuring that the queen is located on one of the frames we transferred them into the observation beehive and fixated their empty sides to the back surface of the hive. Before recording, we allowed each colony to adjust for two weeks after the transfer from the apiary to the observation hive.

To adapt the detection method to the new set of recordings we spatially scaled down the labeled images from D1 and D2 by a factor of 2. The labeled positions were adjusted accordingly, the sizes of foreground regions were equally scaled down to cover similar proportion of bee body surface in the smaller images. We retrained the same approach on the rescaled data and generated predictions of this way trained network on several initial frames of recordings L1-4 and S1-4. To compensate for potential noise introduced by image rescaling and to incorporate images of empty parts of the hives in the training set we used the labelling interface (https://github.com/oist/DenseObjectAnnotation) to correct the predictions in up to five initial frames of each recording. We then retrained the detection model for 10 training iterations on the additional set of labels. This way, with relatively small amount of manual labeling, we adjusted the method to the new recordings reaching comparable accuracy to the one achieved on images on which the method was developed (Supplemental Fig. S1).

During inference, we used windows of size 256 × 256 pixels, which overlapped by a margin of 50 pixels. Any objects in the image margin were discarded.

### Brood Detection

We additionally devised a method for detecting brood cells in a colony. We first used a background extraction method^71,72^ applied to a range of images spanning 12 h. One background image was generated for each consecutive 12 h (Supplemental Movie M1). We then adopted our bee annotation tool (https://github.com/oist/DenseObjectAnnotation) to annotate center points of each brood cell in the generated background images. With the use of this tool we labelled three initial background images from recordings L1-4 (Supplemental Fig. S3).

Based on position labels we generated segmentation labels, in which pixels within circles of radius of 10 pixels around each position label were labeled as foreground. The U-Net segmentation network^12^ was trained to reproduce these segmentation labels. We applied a weighing scheme in which a 2D gaussian scaled by a factor of 10 was generated and centered over each foreground pixel patch. Such weighing was designed to compensate for the class imbalance between the foreground and background pixels and allowed the accurate detection of the centers of each segmentation label.

The network was trained for 1000 epochs with batch size = 16 after which foreground pixel detection accuracy was > 90%, resulting in detection accuracy in the train set of TPR ~ 0.95, FPR ~ 0.02, FNR ~ 0.05. Given the relative ease of the detection task and our extensive previous analysis of the testing accuracy for bee bodies, no further analysis of the brood detection accuracy was performed. Based on manual inspection however, the method generalized to the remaining parts of the recordings as well as to the recording L5 (Supplemental Fig. S2, Supplemental Movie M2).

### Position matching algorithm

For the short-term recordings, we devised a position matching procedure for linking object detections in consecutive video frames into object trajectories. We used both position coordinates as well as the posture categories full-bees and cell-bees. In the first step, all detections are considered as trajectories of length one forming the initial set of assembled trajectories *T* = {*T*_*i*_: *i* = 1. . *n*}. In each following step, detections in consecutive video frames are considered as potential extensions of the trajectories *T*. In step *i*, we calculate the Euclidian distances of the last position of each of the assembled trajectory to detections in the frame at time *t*_*i*_. For a given trajectory *T*_*j*_ with its last position at time point *t*_*j*_ only detections below a given distance cutoff *c*_*d*_ are considered as a match. We define the cutoff as: 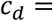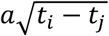 if more than 5 of the last 10 positions in a trajectory are a full-bee and 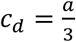 otherwise, where *a* is half of a bee longest dimension, that is 40 px in our recordings. For all detections below this cutoff an additional length factor is added to the Euclidian distance:

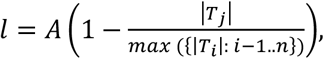

where *A* = 30 is a scaling factor chosen based on accuracy of matching. The length factor prioritizes longer trajectories instead of short trajectories that might arise as an effect of false positive detections.

After all pairs between detections in frame at time *t*_*i*_ and last positions of the assembled trajectories are found and their distances calculated, the matchings are generated in the incremental order of distances until no more matched pairs are found. Matched detections are added to the respective trajectories. Unmatched detections in frame at time *t*_*i*_ are added as potential starting points of new trajectories. Unmatched trajectories are kept until the time between the last detection in that trajectory and the current time is below a predefined gap cutoff. We define the gap cutoff as 10 sec if more than five of the last 10 positions in a trajectory are classified as cell-bee, 1 sec if the last detection in that trajectory is close to the hive entrance, and 3 sec otherwise. This choice of cutoffs is based on the observations that bees inside of the honeycomb cells can be occluded for long timespans and that large densities and fast motion near the entrance can lead to wrong matching if longer gaps are allowed. Assembled trajectories that exceed the gap cutoff are considered as finished and stored for further analysis if their length is above 1 min and discarded otherwise.

We parallelized the matching procedure by splitting the short-term recordings into segments of 1 min. Results of matching within these segments are then matched based on the criteria above. With this parallelization our approach can scale to recordings of arbitrary length with lower computational cost.

### Visual features learning

To improve the accuracy of the trajectory reconstructions, we devised a novel method to exploit the visual features of bee detections via a CNN architecture ^47^ previously shown to perform well in the bee recognition task ^35^, and which we altered by adding a triplet loss in the objective function ^48,49^. The triplet function was originally designed for learning vector embeddings capturing similarity among entities. In the tracking context it is a function where a correct (positive) matching of detections is compared to an incorrect (negative) one. During training, triplets of bee detections are used as input including: (1) anchor image: bee detection in a frame at a time point *t*, (2) positive match: the same bee detection in frame at a time point *t* + Δ*t*, (3) negative match: a different bee in the frame at a time point *t* + Δ*t*. The objective function penalizes representations that set the positive match further apart in terms of the Euclidian distance between the vector embedding of the images (1) and (2) than the distance of the negative match between vector embeddings of the images (1) and (3). To ensure separation of the positive and negative matches a margin *α* is added to the loss:

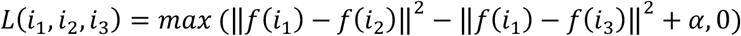

where *i*_1_, *i*_2_, *i*_3_ are the input images (1), (2), (3), *f* is the embedding and the margin *α* = 0.5. The loss value in a training batch is defined as mean loss of all triplets over the number of correct triplets with *L* = 0.

The number of possible triplet bee images in colonies of ~1,000 bees filmed with high temporal resolution prohibits training the network within a reasonable time. Therefore, we implemented two elements in the training procedure to accelerate the learning process. First, the sampling of image triplets is done according to the criteria of the position matching algorithm described above. For a given anchor image only those negative matchings are generated that lie within the time and space distance limit to the anchor image as defined by the criteria of the matching procedure. Corresponding positive matching image is selected from the same video frame as the negative one. Second, in each step of the training, input triplets that show positive value of the triplet loss are fed back into training. All other input triples are randomly sampled according to the rules above.

We tested a range of dimensionalities of the image embeddings (from 16 to 4,096) and chose 64 as the dimensionality offering the best performance in trajectory matching, assessed as a proportion of correctly reconstructed reference trajectories, described below. We also included several data augmentation procedures including random 90° rotations and mirror random flip along both axes. Finally, we implemented background masking and this way trained a model capturing visual features of the bee only, excluding background (Supplemental Fig. S3).

Training comprised 5,000 epochs of 128,000 batches where batch size is 32. After this number of iterations, value of the loss function and the number of incorrect matchings in each batch did not show a decreasing trend. The network was implemented in TensorFlow^73^, trained with Adam optimizer ^74^ using a base learning rate of 0.0001.

### Matching procedure with visual features

Quantitative representations of visual features of bee detections were integrated in the trajectory matching procedure as follows. Trajectories were assembled via analogous sequential video frame processing applying the same time and space distance cutoffs as in the position-based approach described above. For a given trajectory *T*_*j*_ composed of detections {*p*_1_. . *p*_n_}, detections in the following video frame below the distance cutoff *c*_*d*_ to *p*_*n*_ are considered. For each detection *d*_*i*_ the visual similarity to trajectory *T*_*j*_ is quantified as:

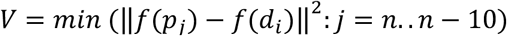

where *f* is vector embedding. Detections with *V* < *c*_*v*_ where *c*_*v*_ = 1.75 is a cutoff for the appearance similarity, are considered as potential extensions of *T*_*j*_.

For all detections below the distance and appearance similarity cutoffs the distance between *T*_*j*_ and a detection *d*_*i*_ is defined as

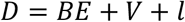

where *E* is the Euclidian distance between last trajectory position *p*_*n*_ and *d*_*i*_, *l* is the length factor as described above, and *B* = 0.033 is a scaling factor chosen based on accuracy of the assembled trajectories. In each step of the matching process matching of trajectories to detections is done in and increasing order of *D*. The matching procedure exploiting visual features follows the same time gap logic and parallelization as the matching based on positions only.

### Training Dataset

To obtain the initial set of trajectories for network training we applied the previously devised “pixel personality”-based method ^35^ on recordings S1-S4 and the results were manually validated. Correct trajectories were then used as the “initial dataset” for training the network on the visual feature embeddings described above.

The trained network was used to infer the vector embeddings of bee detections in videos S1-S4, which we were then used in constructing a new set of trajectories. These trajectories were again manually validated and the correct ones together with the “initial dataset” formed the “final dataset” of trajectories.

The network trained on this way expanded train set was used to derive quantitative representations of bee detections in videos S1-S4 as well as video S5. Images of video S5 were not used in training neither of the detection nor the representation learning network. Trajectories were next constructed based on the visual representations derived via this network and the results were manually validated.

## Acknowledgements

We thank Michael Smith, Dieu My thanh Nguyen and Michael Iuzzolino for comments on the manuscript and code testing. This work was supported by the Okinawa Institute of Science and Technology Graduate University.

## Notes

https://groups.oist.jp/bptu/honeybee-tracking-dataset#tra

## References

1. Seeley, T. D. Honeybee Democracy. (Princeton University Press, 2010).

2. Seeley, T. D. The Wisdom of the Hive: the social physiology of honey bee colonies. (Harvard University Press, 2009).

3. Peleg, O., Peters, J. M., Salcedo, M. K. & Mahadevan, L. Collective mechanical adaptation of honeybee swarms. Nat. Phys. (2018) doi:10.1038/s41567-018-0262-1.

4. Bidari, S., Peleg, O. & Kilpatrick, Z. P. Social inhibition maintains adaptivity and consensus of honeybees foraging in dynamic environments. R Soc Open Sci 6, 191681 (2019).

5. Zayed, A. & Robinson, G. E. Understanding the relationship between brain gene expression and social behavior: lessons from the honey bee. Annu. Rev. Genet. 46, 591–615 (2012).

6. Whitfield, C. W., Cziko, A.-M. & Robinson, G. E. Gene expression profiles in the brain predict behavior in individual honey bees. Science 302, 296–299 (2003).

7. Liang, Z. S. et al. Comparative brain transcriptomic analyses of scouting across distinct behavioural and ecological contexts in honeybees. Proc. Biol. Sci. 281, (2014).

8. Biesmeijer, J. C. & Seeley, T. D. The use of waggle dance information by honey bees throughout their foraging careers. Behav. Ecol. Sociobiol. 59, 133–142 (2005).

9. Wario, F., Wild, B., Couvillon, M. J., Rojas, R. & Landgraf, T. Automatic methods for long-term tracking and the detection and decoding of communication dances in honeybees. Front. Ecol. Evol. 3, (2015).

10. Gernat, T. et al. Automated monitoring of behavior reveals bursty interaction patterns and rapid spreading dynamics in honeybee social networks. Proc. Natl. Acad. Sci. U. S. A. 115, 1433–1438 (2018).

11. Boenisch, F. et al. Tracking all members of a honey bee colony over their lifetime using learned models of correspondence. Frontiers in Robotics and AI 5, 35 (2018).

12. Ronneberger, O., Fischer, P. & Brox, T. U-Net: Convolutional Networks for Biomedical Image Segmentation. in Medical Image Computing and Computer-Assisted Intervention – MICCAI 2015 (eds. Navab, N., Hornegger, J., Wells, W. M. & Frangi, A. F.) 234–241 (Springer International Publishing, 2015).

13. Bozek, K., Hebert, L., Mikheyev, A. S. & Stephens, G. J. Towards dense object tracking in a 2D honeybee hive. in The IEEE Conference on Computer Vision and Pattern Recognition (CVPR) (openaccess.thecvf.com, 2018).

14. Redmon, J., Divvala, S., Girshick, R. & Farhadi, A. You Only Look Once: Unified, Real-Time Object Detection. arXiv [cs.CV] (2015).

15. Newell, A., Yang, K. & Deng, J. Stacked Hourglass Networks for Human Pose Estimation. In Computer Vision – ECCV 2016 483–499 (Springer International Publishing, 2016).

16. Toshev, A. & Szegedy, C. Deeppose: Human pose estimation via deep neural networks. in Proceedings of the IEEE conference on computer vision and pattern recognition 1653–1660 (openaccess.thecvf.com, 2014).

17. Mathis, A. et al. DeepLabCut: markerless pose estimation of user-defined body parts with deep learning. Nat. Neurosci. 21, 1281–1289 (2018).

18. Pereira, T. D. et al. Fast animal pose estimation using deep neural networks. Nat. Methods 16, 117–125 (2019).

19. Fernando, B., Bilen, H., Gavves, E. & Gould, S. Self-supervised video representation learning with odd-one-out networks. in Proceedings of the IEEE conference on computer vision and pattern recognition 3636–3645 (openaccess.thecvf.com, 2017).

20. Wang, X. & Gupta, A. Unsupervised learning of visual representations using videos. in Proceedings of the IEEE International Conference on Computer Vision 2794–2802 (openaccess.thecvf.com, 2015).

21. Taigman, Y., Yang, M., Ranzato, M.’aurelio & Wolf, L. Deepface: Closing the gap to human-level performance in face verification. in Proceedings of the IEEE conference on computer vision and pattern recognition 1701–1708 (cv-foundation.org, 2014).

22. Milan, A., Leal-Taixe, L., Reid, I., Roth, S. & Schindler, K. MOT16: A Benchmark for Multi-Object Tracking. arXiv [cs.CV] (2016).

23. Leal-Taixé, L., Milan, A., Reid, I., Roth, S. & Schindler, K. MOTChallenge 2015: Towards a Benchmark for Multi-Target Tracking. arXiv [cs.CV] (2015).

24. Simonyan, K. & Zisserman, A. Two-Stream Convolutional Networks for Action Recognition in Videos. in Advances in Neural Information Processing Systems 27 (eds. Ghahramani, Z., Welling, M., Cortes, C., Lawrence, N. D. & Weinberger, K. Q.) 568–576 (Curran Associates, Inc., 2014).

25. Carreira, J. & Zisserman, A. Quo vadis, action recognition? a new model and the kinetics dataset. in proceedings of the IEEE Conference on Computer Vision and Pattern Recognition 6299–6308 (openaccess.thecvf.com, 2017).

26. Newby, J. M., Schaefer, A. M., Lee, P. T., Forest, M. G. & Lai, S. K. Convolutional neural networks automate detection for tracking of submicron-scale particles in 2D and 3D. Proc. Natl. Acad. Sci. U. S. A. 115, 9026–9031 (2018).

27. Tsai, H.-F., Gajda, J., Sloan, T. F. W., Rares, A. & Shen, A. Q. Usiigaci: Instance-aware cell tracking in stain-free phase contrast microscopy enabled by machine learning. SoftwareX 9, 230–237 (2019).

28. Liu, K. et al. Fast 3D cell tracking with wide-field fluorescence microscopy through deep learning. arXiv [physics.optics] (2018).

29. Wang, M., Ong, L.-L. S., Dauwels, J. & Asada, H. H. Multicell migration tracking within angiogenic networks by deep learning-based segmentation and augmented Bayesian filtering. J Med Imaging (Bellingham) 5, 024005 (2018).

30. Azevedo, A. W., Gurung, P., Venkatasubramanian, L., Mann, R. & Tuthill, J. C. A size principle for leg motor control in Drosophila. bioRxiv 730218 (2019) doi:10.1101/730218.

31. Bidaye, S. S. et al. Two brain pathways initiate distinct forward walking programs in Drosophila. bioRxiv 798439 (2019) doi:10.1101/798439.

32. Clifton, G. T., Holway, D. & Gravish, N. Rough substrates constrain walking speed in ants through modulation of stride frequency and not stride length. bioRxiv 731380 (2019) doi:10.1101/731380.

33. Romero-Ferrero, F., Bergomi, M. G., Hinz, R., Heras, F. J. H. & de Polavieja, G. G. idtracker.ai: Tracking all individuals in large collectives of unmarked animals. arXiv [cs.CV] (2018).

34. Pérez-Escudero, A., Vicente-Page, J., Hinz, R. C., Arganda, S. & de Polavieja, G. G. idTracker: tracking individuals in a group by automatic identification of unmarked animals. Nat. Methods 11, 743–748 (2014).

35. Bozek, K., Hebert, L., Mikheyev, A. S. & Stephens, G. J. Pixel personality for dense object tracking in a 2D honeybee hive. arXiv [cs.CV] (2018).

36. Tschinkel, W. R. Insect sociometry, a field in search of data. Insectes Soc. 38, 77–82 (1991).

37. Smith, M. L., Ostwald, M. M. & Seeley, T. D. Honey bee sociometry: tracking honey bee colonies and their nest contents from colony founding until death. Insectes Soc. 63, 553–563 (2016).

38. Smith, M. L., Koenig, P. A. & Peters, J. M. The cues of colony size: how honey bees sense that their colony is large enough to begin to invest in reproduction. J. Exp. Biol. 220, 1597–1605 (2017).

39. Lee, K. V., Goblirsch, M., McDermott, E., Tarpy, D. R. & Spivak, M. Is the Brood Pattern within a Honey Bee Colony a Reliable Indicator of Queen Quality? Insects 10, (2019).

40. Klein, B. A., Olzsowy, K. M., Klein, A., Saunders, K. M. & Seeley, T. D. Caste-dependent sleep of worker honey bees. J. Exp. Biol. 211, 3028–3040 (2008).

41. Ostwald, M. M., Smith, M. L. & Seeley, T. D. The behavioral regulation of thirst, water collection and water storage in honey bee colonies. J. Exp. Biol. 219, 2156–2165 (2016).

42. Oldroyd, B. P. What’s killing American honey bees? PLoS Biol. 5, e168 (2007).

43. Magal, P., Webb, G. F. & Wu, Y. An Environmental Model of Honey Bee Colony Collapse Due to Pesticide Contamination. Bull. Math. Biol. (2019) doi:10.1007/s11538-019-00662-5.

44. Dainat, B., Evans, J. D., Chen, Y. P., Gauthier, L. & Neumann, P. Predictive markers of honey bee colony collapse. PLoS One 7, e32151 (2012).

45. Dennis, B. & Kemp, W. P. How Hives Collapse: Allee Effects, Ecological Resilience, and the Honey Bee. PLoS One 11, e0150055 (2016).

46. Romero-Ferrero, F., Bergomi, M. G., Hinz, R. C., Heras, F. J. H. & de Polavieja, G. G. idtracker.ai: tracking all individuals in small or large collectives of unmarked animals. Nat. Methods 16, 179–182 (2019).

47. Szegedy, C., Vanhoucke, V., Ioffe, S., Shlens, J. & Wojna, Z. Rethinking the Inception Architecture for Computer Vision. arXiv [cs.CV] (2015).

48. Chechik, G. Large Scale Online Learning of Image Similarity Through Ranking. J. Mach. Learn. Res. 11, 1109–1135 (2010).

49. Schroff, F., Kalenichenko, D. & Philbin, J. FaceNet: A unified embedding for face recognition and clustering. in 2015 IEEE Conference on Computer Vision and Pattern Recognition (CVPR) 815–823 (2015).

50. Tautz, J. & Lindauer, M. Honeybees establish specific sites on the comb for their waggle dances. Journal of Comparative Physiology A 180, 537–539 (1997).

51. Johnson, B. R. Global information sampling in the honey bee. Naturwissenschaften 95, 523–530 (2008).

52. Graving, J. M. et al. DeepPoseKit, a software toolkit for fast and robust animal pose estimation using deep learning. Elife 8, (2019).

53. Hinz, R. C. & de Polavieja, G. G. Ontogeny of collective behavior reveals a simple attraction rule. Proc. Natl. Acad. Sci. U. S. A. 114, 2295–2300 (2017).

54. Cavagna, A. et al. Scale-free correlations in starling flocks. Proc. Natl. Acad. Sci. U. S. A. 107, 11865–11870 (2010).

55. Meikle, W. G. et al. Using within-day hive weight changes to measure environmental effects on honey bee colonies. PLoS One 13, e0197589 (2018).

56. Marchal, P. et al. Automated monitoring of bee behaviour using connected hives: Towards a computational apidology. Apidologie (2019) doi:10.1007/s13592-019-00714-8.

57. Crall, J. D. et al. Neonicotinoid exposure disrupts bumblebee nest behavior, social networks, and thermoregulation. Science 362, 683–686 (2018).

58. Milan, A., Rezatofighi, S. H., Dick, A., Reid, I. & Schindler, K. Online Multi-Target Tracking Using Recurrent Neural Networks. arXiv [cs.CV] (2016).

59. Ning, G. et al. Spatially Supervised Recurrent Convolutional Neural Networks for Visual Object Tracking. arXiv [cs.CV] (2016).

60. Nixon, H. L. & Ribbands, C. R. Food transmission within the honeybee community. Proc. R. Soc. Lond. B Biol. Sci. 140, 43–50 (1952).

61. Peters, J. M., Gravish, N. & Combes, S. A. Wings as impellers: honey bees co-opt flight system to induce nest ventilation and disperse pheromones. J. Exp. Biol. 220, 2203–2209 (2017).

62. Rivière, J. et al. Toward a Complete Agent-Based Model of a Honeybee Colony. in Highlights of Practical Applications of Agents, Multi-Agent Systems, and Complexity: The PAAMS Collection 493–505 (Springer International Publishing, 2018).

63. Becher, M. A., Osborne, J. L., Thorbek, P., Kennedy, P. J. & Grimm, V. Towards a systems approach for understanding honeybee decline: a stocktaking and synthesis of existing models. J. Appl. Ecol. 50, 868–880 (2013).

64. Torres, D. J., Ricoy, U. M. & Roybal, S. Modeling Honey Bee Populations. PLoS One 10, e0130966 (2015).

65. Betti, M. I., Wahl, L. M. & Zamir, M. Age structure is critical to the population dynamics and survival of honeybee colonies. R Soc Open Sci 3, 160444 (2016).

66. Okada, R. et al. Error in the honeybee waggle dance improves foraging flexibility. Sci. Rep. 4, 4175 (2014).

67. Chen, X., Randi, F., Leifer, A. M. & Bialek, W. Searching for collective behavior in a small brain. Phys Rev E 99, 052418 (2019).

68. Cavagna, A., Giardina, I., Mora, T. & Walczak, A. M. Physical constraints in biological collective behaviour. Current Opinion in Systems Biology 9, 49–54 (2018).

69. Mersch, D. P., Crespi, A. & Keller, L. Tracking individuals shows spatial fidelity is a key regulator of ant social organization. Science 340, 1090–1093 (2013).

70. Bain, N. & Bartolo, D. Dynamic response and hydrodynamics of polarized crowds. Science 363, 46–49 (2019).

71. Laugraud, B., Piérard, S., Braham, M. & Van Droogenbroeck, M. Simple Median-Based Method for Stationary Background Generation Using Background Subtraction Algorithms. in New Trends in Image Analysis and Processing -- ICIAP 2015 Workshops 477–484 (Springer International Publishing, 2015).

72. Laugraud, B., Piérard, S. & Van Droogenbroeck, M. LaBGen: A method based on motion detection for generating the background of a scene. Pattern Recognit. Lett. 96, 12–21 (2017).

73. Abadi, M. et al. TensorFlow: Large-Scale Machine Learning on Heterogeneous Distributed Systems. arXiv [cs.DC] (2016).

74. Kingma, D. & Ba, J. Adam: A Method for Stochastic Optimization. arXiv [cs.LG] (2014).

75. Maaten, L. van der & Hinton, G. Visualizing Data using t-SNE. J. Mach. Learn. Res. 9, 2579–2605 (2008).

