## Supplemental Material for "Markerless tracking of an entire insect colony"

#### 1. Figures

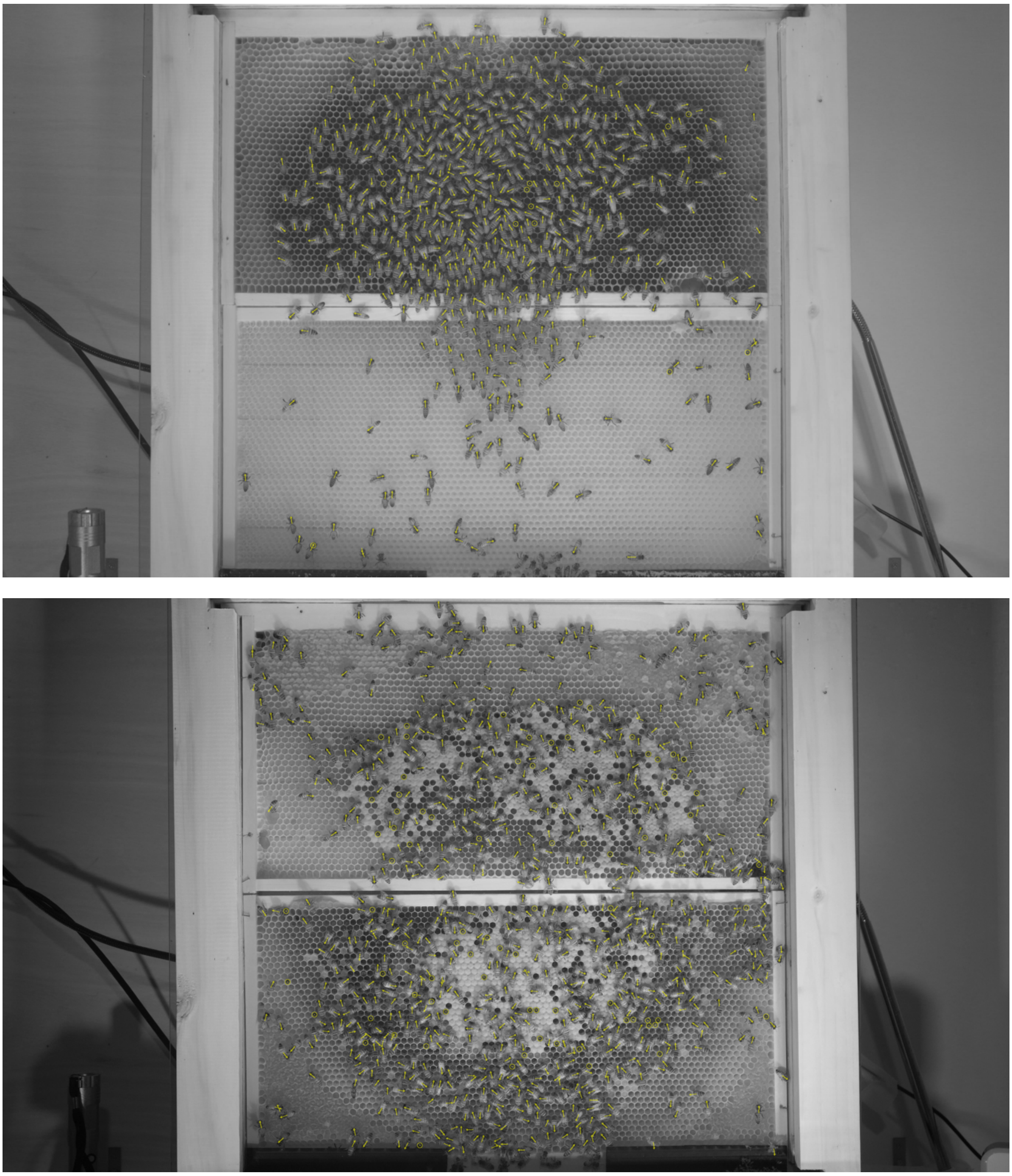

S1. Example predictions from the frames in middle of recordings L5 (upper panel) and S5 (bottom panel). Each detections' center is marked with a dot. Detections recognized as cell-bees are marked with round symbols, full-bees are marked with arrows that indicate the bee head-tail orientation angle. These two recordings were not part of the training set for the detection and tracking methods.

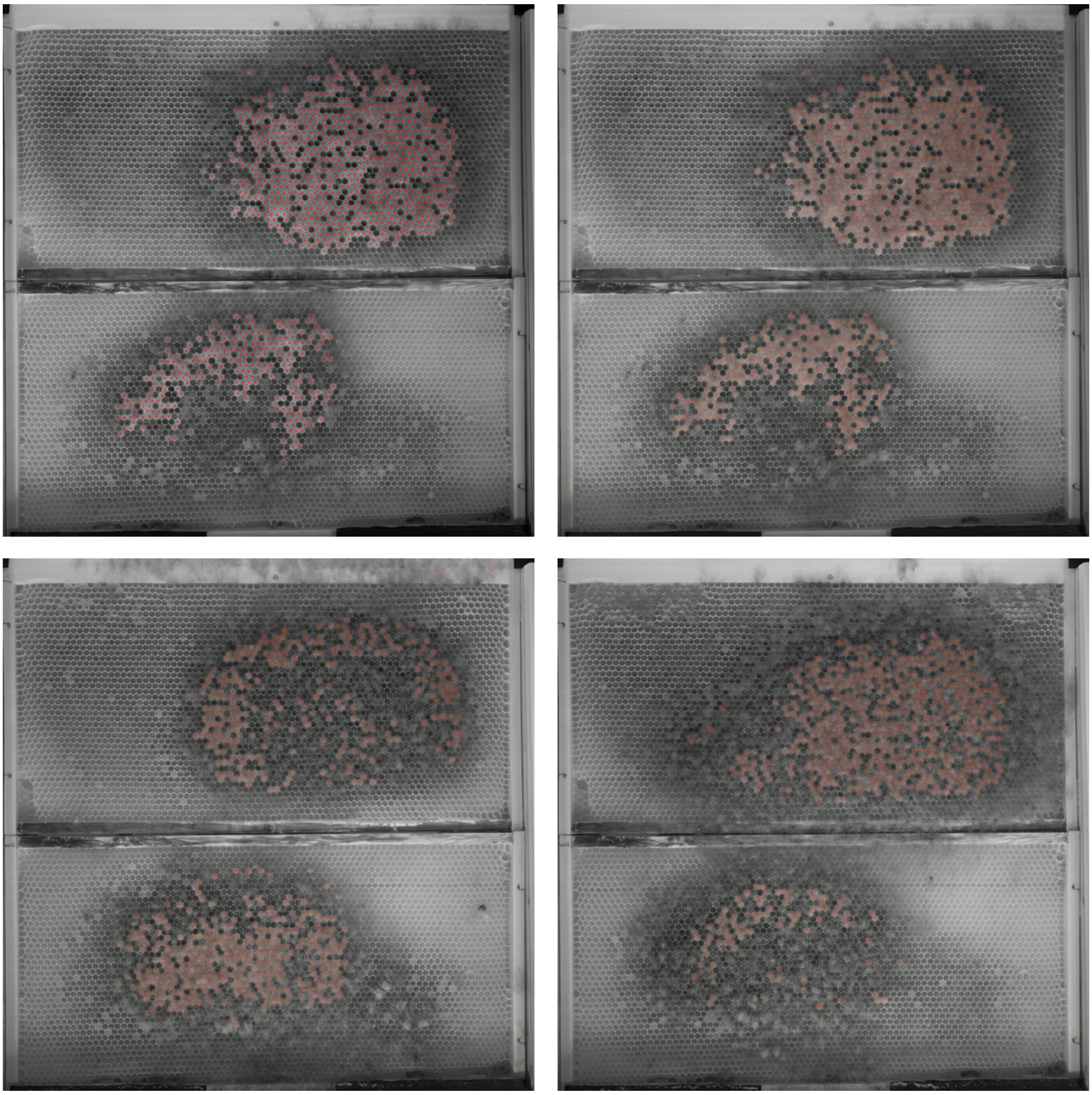

S2. Example of brood cell labeling and predictions in video L2. Labeling and network prediction of the brood image of the first 12 h of the recording are shown in the upper left and right panels respectively. Predictions of brood in the following frames 10 and 20 days later, respectively are shown in the bottom panels.

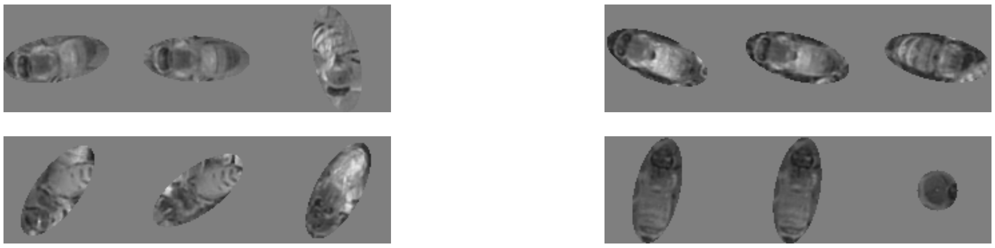

S3. Masking of background. To investigate the role of background in the accuracy of the matching procedure ellipse- and round-shaped masks were applied on the bee images during training and testing. Ellipse-shaped masks were applied to the full-bee images and rotated according to the bee body axis. Round-shaped masks were applied to the cell-bees. The images represent triplets of images that are fed into the network during training. From left to right each image contains anchor image, positive match, and negative match.

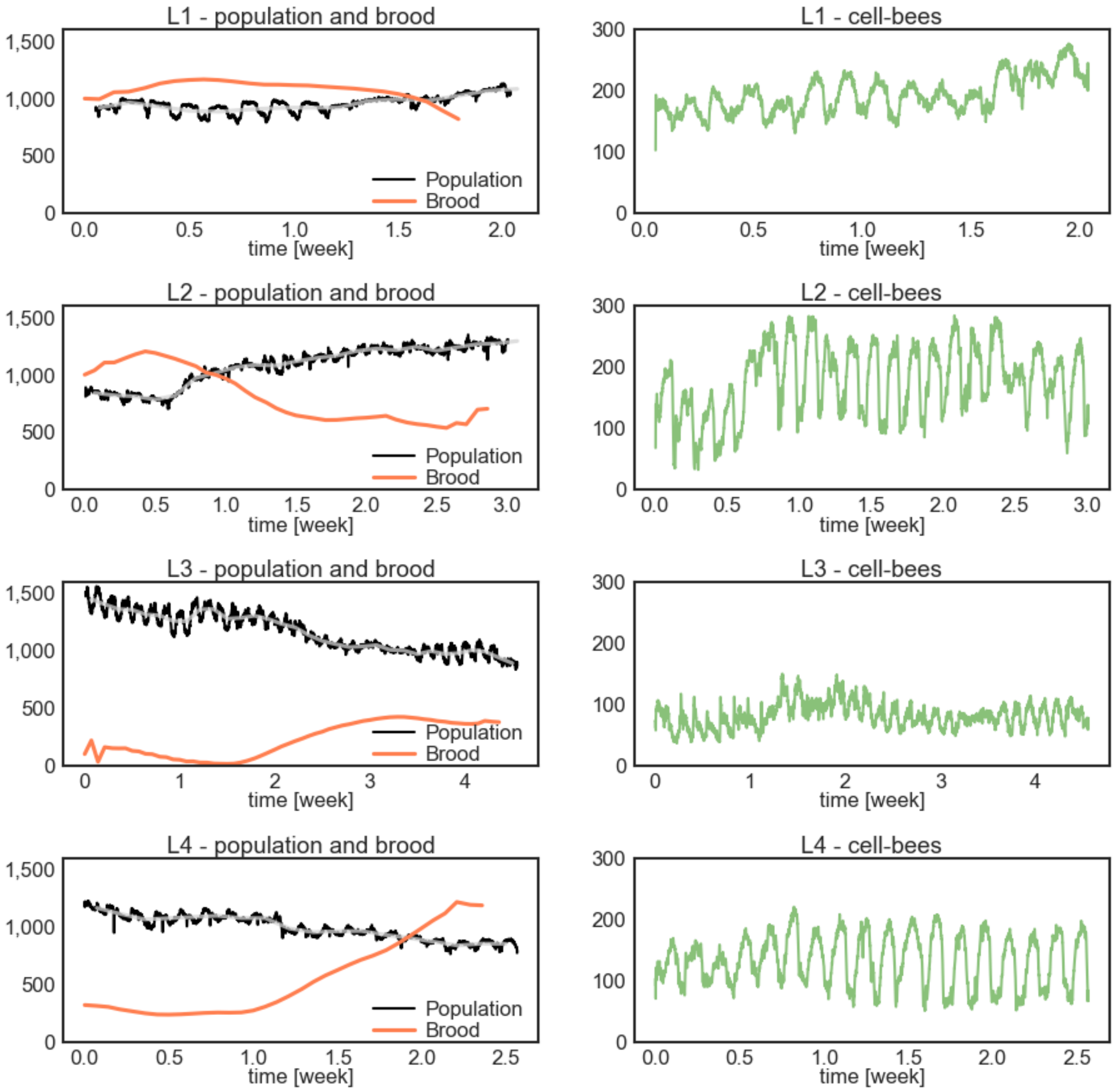


S4. Population, brood cell, and cell-bee counts in recordings L1-L4.

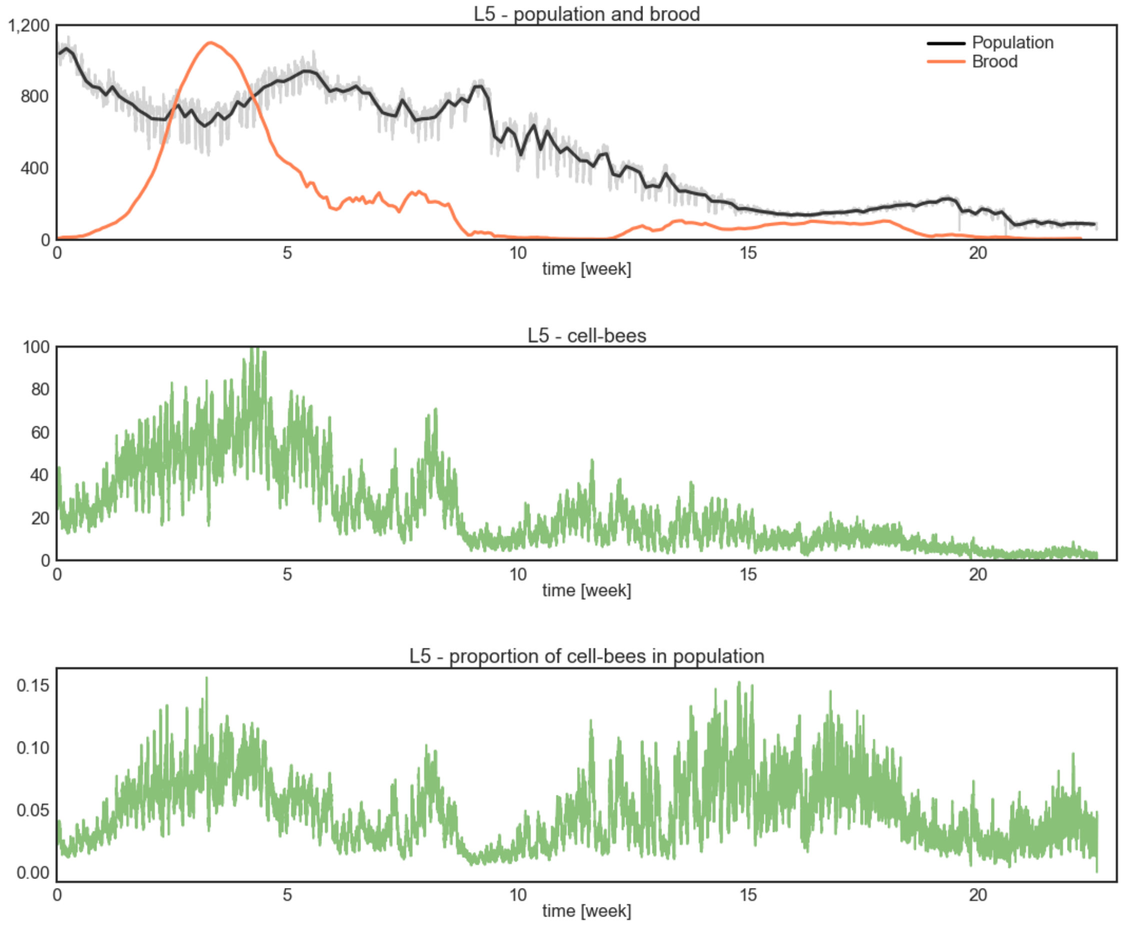


S5. Population, brood cell, and cell-bee counts in beehive L5. Bottom panel shows numbers of cell-bees as a proportion of all bees detected in the hive in a given frame of the recording. Regardless of normalization method, the daily fluctuation of this count is present.

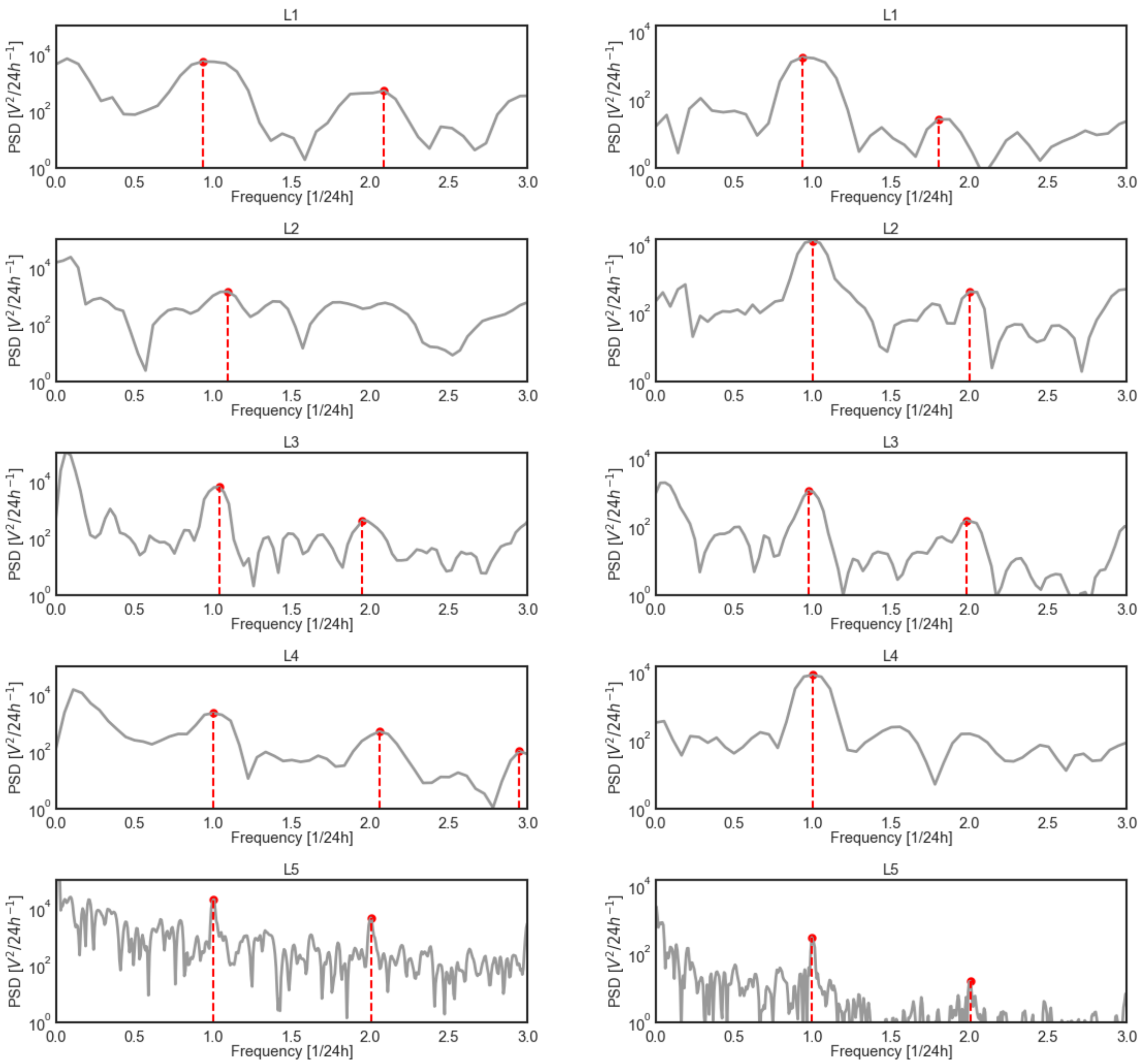


S6. Power spectrum distribution of the population change (left panel) and cell-bee count (right panel), in recordings L1-L5. Local maxima, calculated over 5 h timespans are indicated with red dots in each plot.

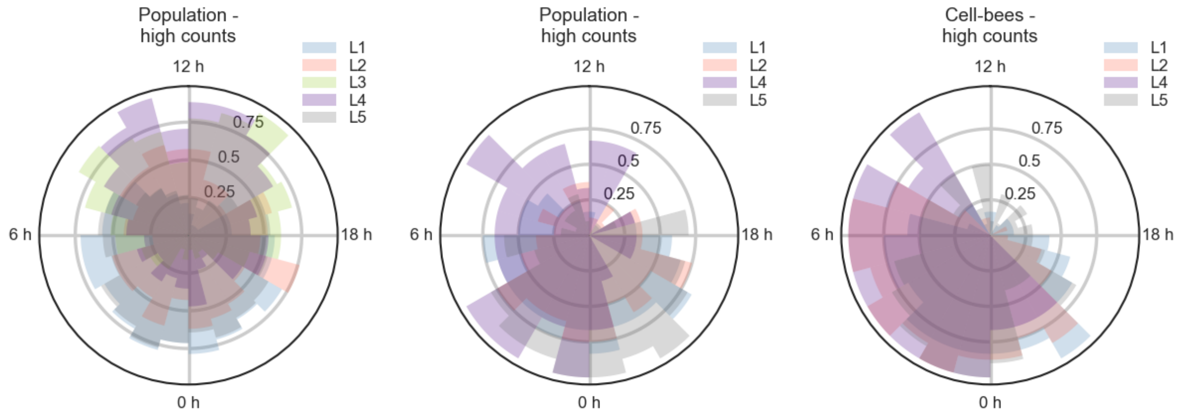


S7. Distributions of hours when the highest population (left and middle panel) and cell-bee numbers (right plot) are present in hives L1-L5. Analogous to Fig. 3E, circular plots are separated into 24 bins. For each 24 h of a recording the number of times that the population or cell-bees counts are above the median number in the respective 24 h time window is counted for each 1 h bin and averaged over the number of days in the recording. Middle and right plots contain only days when number of brood cells in these hives exceeds 800. While no clear phase pattern of the population fluctuation among the hives is present (left plot), on days with high brood count high numbers of bees are present predominantly at night which might reflect foraging activity in these hives (middle plot). On the same days the clear phase of cell-bees (right plot) is more pronounced compared to all recording days shown in Fig. 3E.

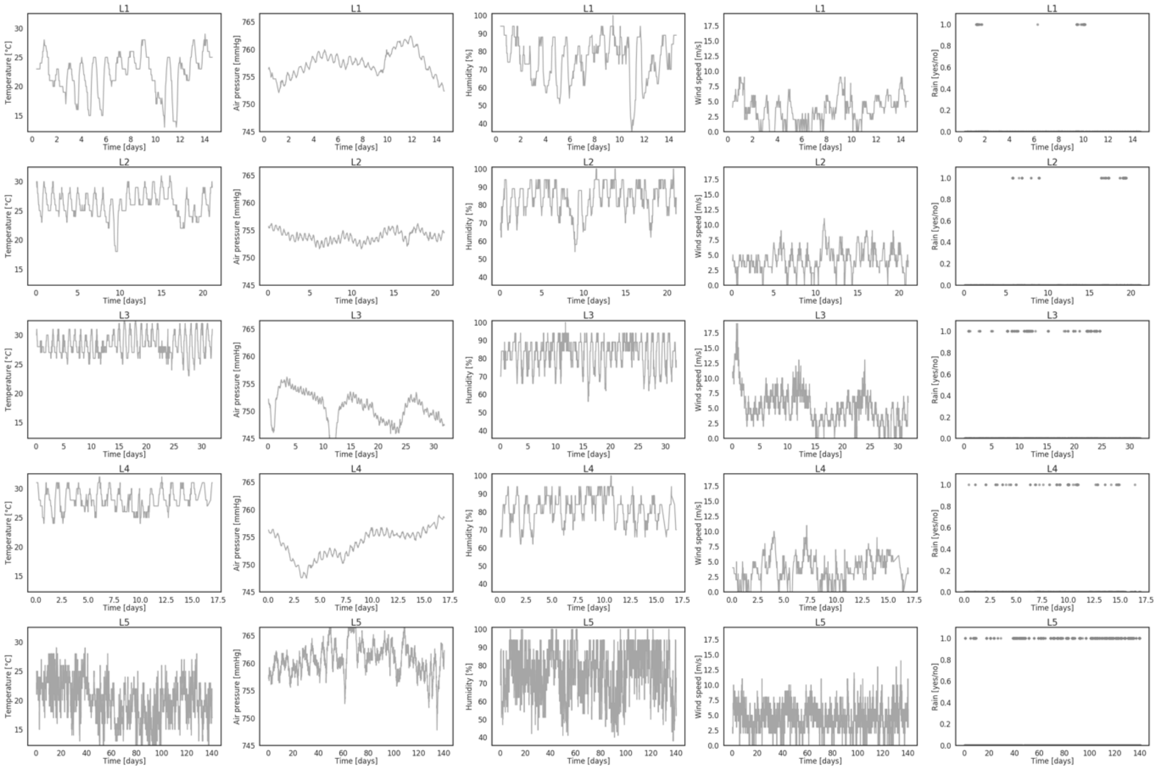

S8. Weather conditions during the time of recordings L1-L5. Shown are temperature, air pressure, humidity, wind, and rain reported in the location of the hives. While no extreme weather events were present during the recordings, hives L3 and L4 were recorded during particularly high temperatures which might be the reason for the anticorrelated phase of the population and cell-bee count.

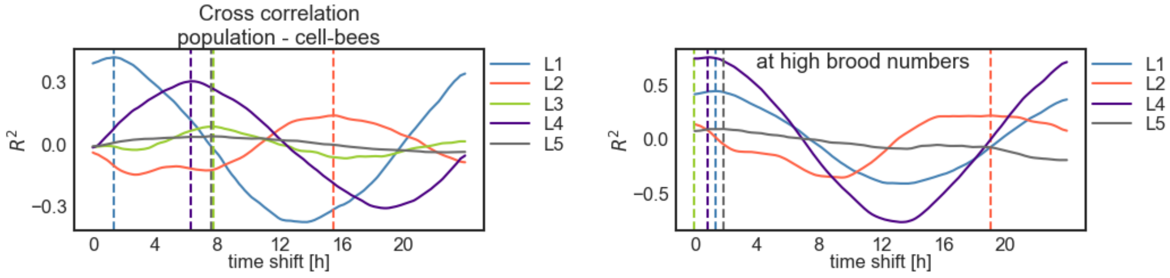


S9. Temporal cross-correlation between the total and cell-bee populations. Correlations were calculated in temporal shifts of 2 min and we indicate the shift with maximum correlation by a vertical dashed line. Except for recording L1, the total population and number of cell-bees show an approximately 8 h phase shift. For L3 and L4 the shift of the total population is ~8 h forward relative to the bell-bee population, while for L2 the shift is equivalently ~8 h backward or ~16 h forward. The bottom plot shows an analogous cross correlation analysis calculated on days of the recordings when the brood numbers are above 800. On those days the total and cell-bee populations are more closely in phase.

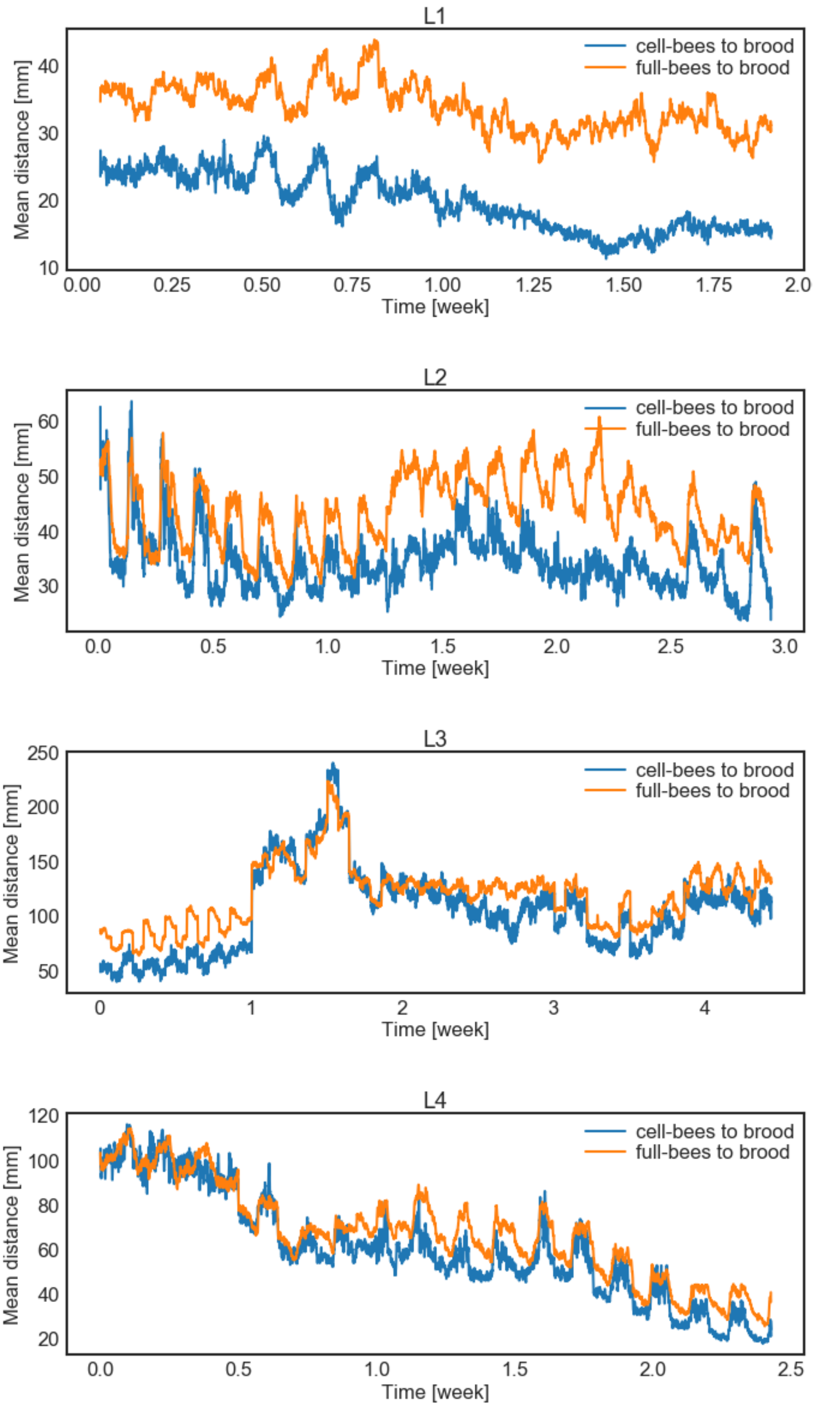


S10. Mean distance of each full-bee (blue) and each cell-bee (orange) to three closest brood cells. Both distances show daily fluctuations with peak times during the day suggesting that at night bees regroup around the brood. Bees inside comb cells tend to be closer to the brood cells suggesting the role of this activity in brood-related activities, such as thermoregulation.

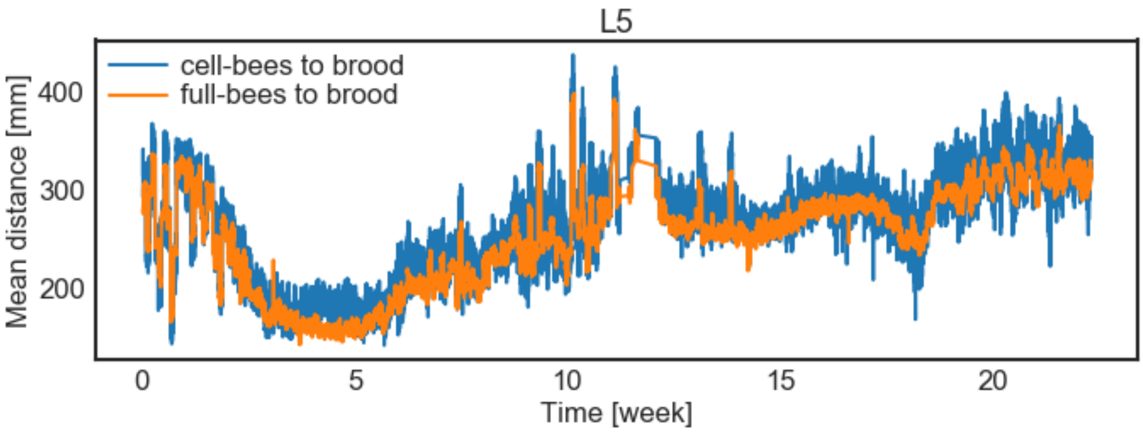


S11. Analogous to S9 in recording L5.

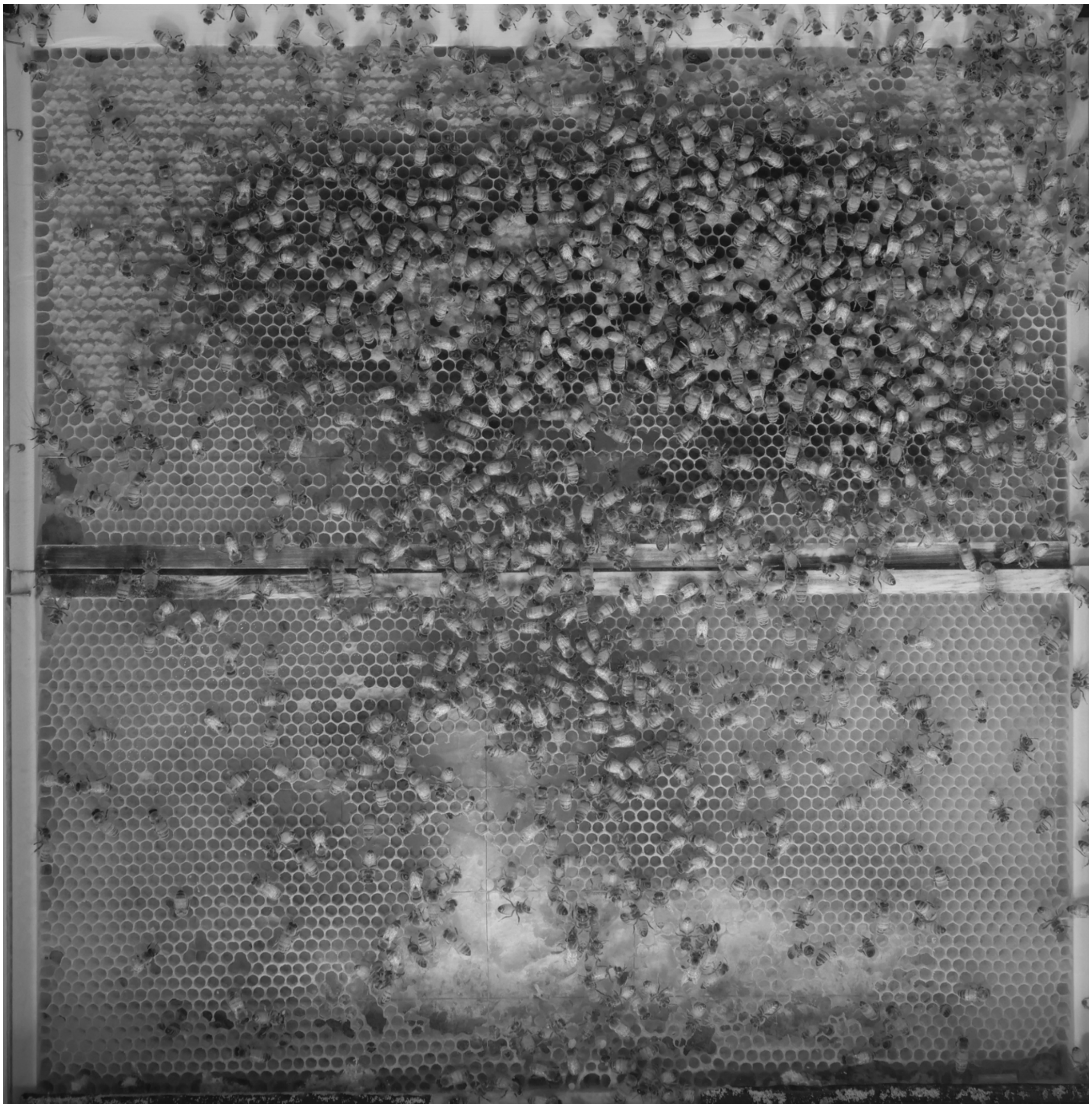

S12. Image of the hive L3 in the third week of the recording. Bottom frame was damaged due to moth infestation.

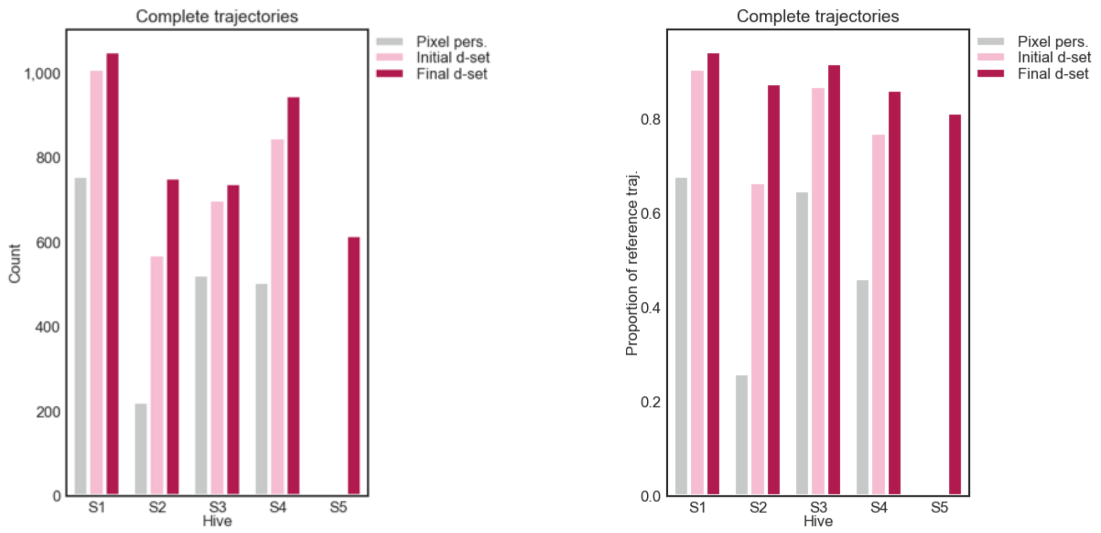


S13. Analogous to Fig. 3C – complete trajectories constructed with the respective methods and quantified as counts (left panel) and proportion of the number of all reference trajectories collected in beehives S1-S5.

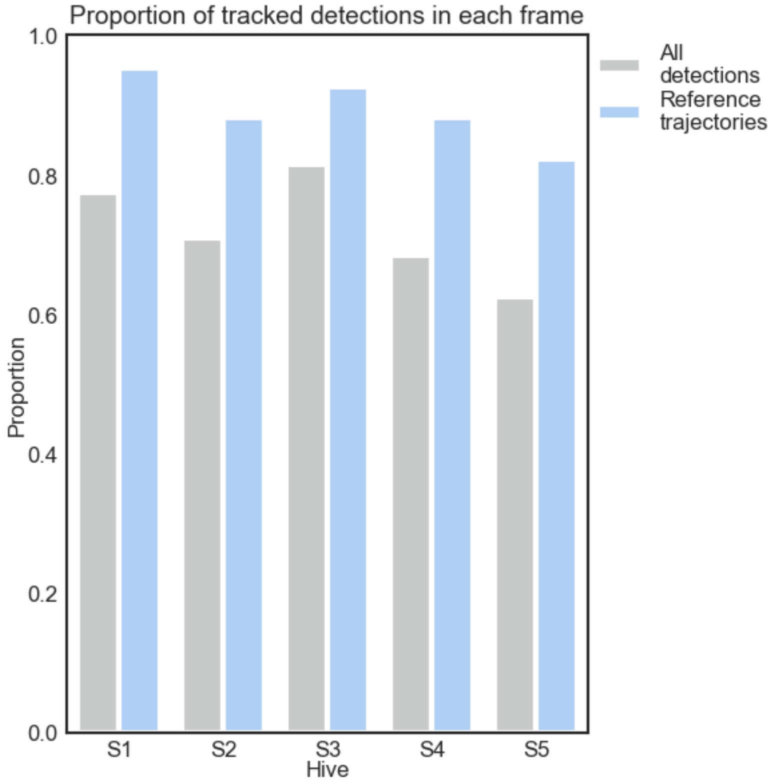


S14. Mean proportion of detections in each frame that belong to a correct trajectory relative to all detections in a frame (gray) and relative to detections that belong to any reference trajectory (blue). The trajectories included in this plot were constructed based on the 'final dataset' model.

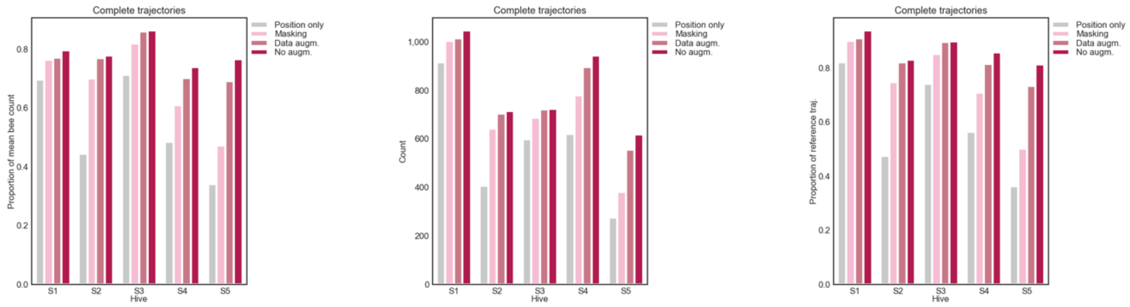


S15. Comparison of the number of correct trajectories constructed without and with augmentation and masking procedures. The numbers shown are relative to average number of detections in each respective hive (left), raw counts (middle), and relative to the number of reference trajectories in the respective hives (right). Masking and augmentation appear not to improve tracking results.

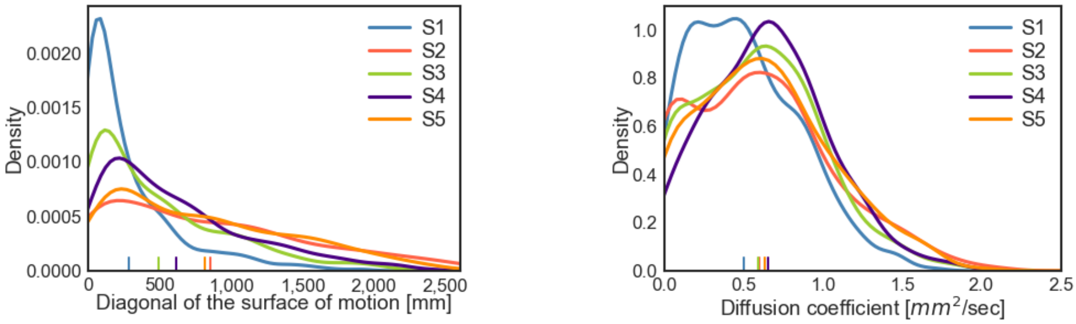


S16. Analogous to Fig. 5A-B, distributions of trajectory surface (left) and diffusion coefficient (right) in hives S1-S5. While differences in motion across the hives is observable based on the trajectory surface, mean diffusivity of the hives is comparable.

### 2. Tables

T1. Accuracy of detection.

| | TP | FP | Error: | | | |
|---|---|---|---|---|---|---|
| | | | Class: | Position [pixel] | Axis [°] | Orientation angle [°] |
| **Human labeling** | - | 0.07 | 0.04 | 6.7 | - | 7.7 |
| **Our method** | 0.96 | 0.06 | 0.19 | 5.1 | 8.8 | 9.7 |

T2. List of recordings. Beehives were imaged in two different locations with varying camera resolutions. All imaging data in this study was collected in 2019.

| Recording | Location | Start date and time | Pixel resolution | Time resolution | Number of frames |
|---|---|---|---|---|---|
| L1 | 2 | 23-04 20:55 | 2560 x 2560 | 1 / 2 min | 9,999 |
| L2 | 2 | 15-05 13:00 | 2560 x 2560 | 1 / 2 min | 15,124 |
| L3 | 2 | 09-07 14:30 | 2560 x 2560 | 1 / 2 min | 22,892 |
| L4 | 2 | 31-09 12:00 | 2560 x 2560 | 1 / 2 min | 12,946 |
| L5 | 2 | 26-10 17:00 | 3840×2160 | 1 / 1 min | 227,131 |
| S1 | 1 | 12-12 09:40 | 5120x5120 | 30 fps | 9,000 |
| S2 | 1 | 15-05 10:00 | 5120x5120 | 30 fps | 9,000 |
| S3 | 2 | 27-11 10:00 | 3840×2160 | 30 fps | 9,000 |
| S4 | 2 | 25-02 11:30 | 3840×2160 | 30 fps | 9,000 |
| S5 | 2 | 19-09 09:30 | 3840×2160 | 30 fps | 9,000 |

T3. Completeness of the tracking results. The table lists the number of correct trajectories obtained with the presented methods in recordings S1-S5. Column 'mean bee count' lists the average number of detection in each video frame. 'Reference trajectories' lists the total number of trajectories that were collected and validated as correct for the respective beehives. The trajectories constructed by each tracking methods are matched against these reference trajectories. Following columns list the raw number of correct trajectories found by each method and the same number as a proportion to the mean number of detections in parentheses. Results of the best performing method are marked in red.

| **Recording** | **Mean bee count** | **Reference trajectories** | **Position only** | **Pixel personality** | **Initial d-set** | **Final d-set** | **Data augmentation (final d-set)** | **Background masking (final d-set)** |
|---|---|---|---|---|---|---|---|---|
| S1 | 1315.6 | 1115 | 916 (0.696) | 758 (0.576) | 1010 (0.768) | 1046 (0.795) | 1014 (0.771) | 1004 (0.763) |
| S2 | 917.4 | 859 | 408 (0.445) | 223 (0.243) | 571 (0.622) | 714 (0.778) | 705 (0.769) | 642 (0.700) |
| S3 | 839.1 | 806 | 598 (0.713) | 523 (0.623) | 700 (0.834) | 724 (0.863) | 722 (0.860) | 687 (0.819) |
| S4 | 1278.3 | 1100 | 620 (0.485) | 507 (0.397) | 848 (0.663) | 944 (0.738) | 896 (0.701) | 779 (0.609) |
| S5 | 805.5 | 758 | 275 (0.341) | - | - | 617 (0.766) | 556 (0.690) | 381 (0.473) |

### 3. Movie Legends

Supplemental movies can be found under the address: https://groups.oist.jp/bptu/honeybee-tracking-dataset#tra

M1. Background extracted from hive L1. Each frame represents 12 h of the original recording.

M2. Background extracted from hive L5. Each frame represents 12 h of the original recording, predicted brood cells are marked in red.

M3-M7. Example bee trajectories with their corresponding trajectories through the space of visual features. Examples shown are from hives S1-S5, respectively. The originally 64-dimensional vectors of visual features representing each bee instance are projected into 3D with the use of t-Distributed Stochastic Neighbor Embedding (t-SNE). Analogous to the plot 3B, 10 most recent representations of the tracked bee are marked in red dots and representations of all other bees from three most recent video frames are marked in yellow dots.

M8-M12. All reference trajectories in hives S1-S5. The short video snippets are provided to visually illustrate the proportion of tracked individuals in each hives.

M13-M15. Bees performing waggle dance, whose trajectories are shown in Fig. 4D.

M16-M18. Bees visiting multiple comb cells, whose trajectories are shown in Fig. 4F.
